# Single-cell quantitative bioimaging of *P. berghei* liver stage translation

**DOI:** 10.1101/2023.07.05.547872

**Authors:** James L. McLellan, William Sausman, Ashley B. Reers, Evelien M. Bunnik, Kirsten K. Hanson

## Abstract

*Plasmodium* parasite resistance to existing antimalarial drugs poses a devastating threat to the lives of many who depend on their efficacy. New antimalarial drugs and novel drug targets are in critical need, along with novel assays to accelerate their identification. Given the essentiality of protein synthesis throughout the complex parasite lifecycle, translation inhibitors are a promising drug class, capable of targeting the disease-causing blood stage of infection, as well as the asymptomatic liver stage, a crucial target for prophylaxis. To identify compounds capable of inhibiting liver stage parasite translation, we developed an assay to visualize and quantify translation in the *P. berghei-*HepG2 infection model. After labeling infected monolayers with o-propargyl puromycin (OPP), a functionalized analog of puromycin permitting subsequent bioorthogonal addition of a fluorophore to each OPP-terminated nascent polypetide, we use automated confocal feedback microscopy followed by batch image segmentation and feature extraction to visualize and quantify the nascent proteome in individual *P. berghei* liver stage parasites and host cells simultaneously. After validation, we demonstrate specific, concentration-dependent liver stage translation inhibition by both parasite-selective and pan-eukaryotic active compounds, and further show that acute pre-treatment and competition modes of the OPP assay can distinguish between direct and indirect translation inhibitors. We identify a Malaria Box compound, MMV019266, as a direct translation inhibitor in *P. berghei* liver stages and confirm this potential mode of action in *P. falciparum* asexual blood stages.

## Introduction

*Plasmodium* parasites are the causative agent of malaria and continue to have an outsized effect on global public health, causing an estimated 241 million cases in 2020, with 77% of deaths occurring in children under the age of five (1). Antimalarial drugs are essential for treating malaria, however, all currently used antimalarials are associated with parasite resistance. The spread of *kelch13-*mediated resistance to the front-line antimalarial artemisinin in South East Asia and its recent *de novo* emergence in Rwanda demonstrate the critical threats to the efficacy of artemisinin combination therapies (ACTs), the front-line therapeutics targeting asexual blood stage (ABS) parasites, which cause all malaria symptoms (2–5). In addition to treating malaria, antimalarial drugs would ideally be able to clear any non-replicative gametocytes in the blood, preventing transmission back to the mosquito vector. Antimalarials are also crucial for disease prophylaxis, with the *Plasmodium* liver stage a key target to prevent both disease and transmission (6). Attractive antimalarials would thus have activity against each of these 3 stages despite significant stage-specific differences in biology (7–9), highlighting the utility of targeting core cellular processes, like translation, that are crucial for all mammalian stages of development.

Translation of mRNA nucleotide sequences to amino acids during the ribosomal synthesis of proteins is a central evolutionarily conserved cellular process that has been extensively targeted with antibiotics treating bacterial infections (10), but *Plasmodium* translation has not been targeted by any clinically approved antimalarials to date. *Plasmodium* protein synthesis is a highly desirable process to target, as translation can be blocked via many different molecular targets. DDD107498 (also known as cabamiquine and M5717), which is thought to target eEF2, a core component of polypeptide elongation on the ribosome (11), and a number of cytoplasmic aminoacyl-tRNA synthetase (aaRS) inhibitors, which prevent the linkage between a tRNA and its cognate amino acid, are in various stages of clinical and pre-clinical development, respectively (12, 13). Additionally, many pan-eukaryotic translation inhibitors have antiplasmodial activity against *P. falciparum* (Pf) ABS in standard 48 hour (h) assays, and were shown to directly target the Pf cytoplasmic translation apparatus using a bulk ABS lysate approach in which translation of the exogenous luciferase transcript is used as a biomarker for total cellular translation (14, 15). Currently, the ability to gain such mechanistic information about antimalarial activity is almost entirely dependent on ABS experiments (16), with the assumption that antiparasitic activity in other stages occurs via the same mechanism. Liver stages are particularly problematic as they rely on highly metabolically active hepatocytes for their own development, which makes bulk population readout of conserved processes like translation impossible due to the signal from hepatocytes themselves. It also complicates interpretation of liver stage antiplasmodial activity, as it may integrate both hepatocyte- and parasite-directed effects.

Here, we report a bioimage-based assay quantifying *P. berghei* liver stage translation in the native cellular context. We rely on the activity of the aminoacyl-tRNA mimic puromycin, which is covalently bound to the C-terminus of a nascent polypeptide during the elongation reaction, causing the ribosome to disassociate and release the puromycin-bound nascent-polypeptide (17–21). A synthetic puromycin analog, puromycin (OPP), was shown to truncate and label nascent polypeptides in an identical manner, but contains a small alkyne tag, facilitating the copper-catalyzed cycloaddition of a picolyl azide fluorophore in a bioorthogonal reaction, commonly termed “click chemistry” (22). Combining the OPP labeling of nascent polypeptides with automated fluorescence microscopy and quantitative image analysis, we demonstrate specific and separable *in cellulo* quantification of *P. berghei* and *H. sapiens* translation during liver stage development in HepG2 cells, and use the assay to identify both direct and indirect inhibitors of *Plasmodium* liver stage translation.

## Results

### Visualization of the *Plasmodium* nascent proteome

With a goal of quantifying translation in single parasites, we first explored whether OPP would label the *P. berghei* nascent proteome during liver stage development. Infected HepG2 cells were treated with OPP for 30 minutes at 37°C, then immediately fixed with 4% paraformaldehyde, which stops the labeling reaction and preserves the quantity and cellular localization of the OPP-labeled polypeptides (22). Post-fixation, a click chemistry reaction attaches a picolyl azide conjugated fluorophore to the OPP-labelled polypeptides of both host and parasite, which can then be visualized with fluorescence microscopy. AlexaFluor555 was used to visualize newly synthesized peptides throughout this study, and the resulting signal, from both host and parasite nascent proteomes, will be referred to as OPP-A555. As expected, we can visualize translation throughout liver stage parasite development (Fig. 1A), from newly invaded sporozoite (2 hpi), through merozoite formation (57 hpi). By eye, parasite translation intensity (evidenced by OPP-A555 signal) appears generally greater than that of the host cell and surrounding non-infected HepG2 cells. The robust and highly specific OPP-A555 signal (Fig. S1) suggests that this approach can be adapted to directly quantify translation of the intrahepatic parasite. The OPP labeling technology is particularly flexible, as it does not require any genetic modifications to label the nascent proteome and should thus be directly adaptable to a wide variety of organisms, including other *Plasmodium* species and stages. Supporting this, *P. falciparum* asexual blood stage translation can also be visualized in infected erythrocytes using a highly similar protocol (Fig. S2).

**Figure 1.**
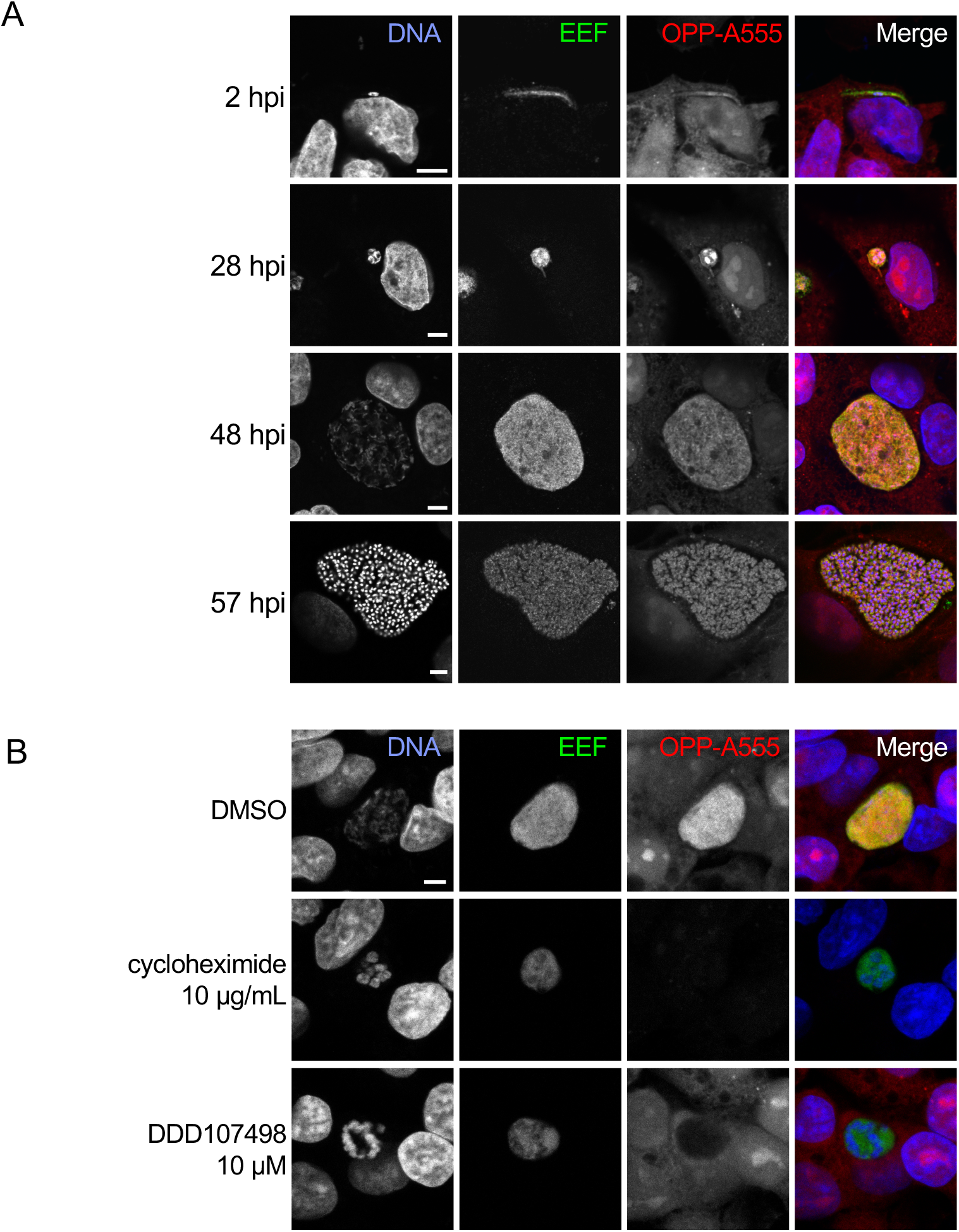
Visualization of the nascent proteome in *Plasmodium berghei* liver stage parasites. A-B) Representative, single confocal images of *P. berghei-*infected HepG2 cells, with OPP conjugated to Alexa Fluor 555 (OPP-A555) labeling the nascent proteome in both HepG2 and parasite (EEF), with Hoechst labeling DNA. Single channel images are shown in grayscale, with merges pseudo colored as labeled. A) Visualization of the nascent proteome throughout liver stage development, with parasite immunolabeled with α-UIS4 (2 hpi) or α- HSP70 (28, 48, and 57 hpi). B) Nascent proteome visualized in infected HepG2 cells following treatment from 44-48 hpi with translation inhibitors cycloheximide (10 µg/mL), or DDD107498 (10 µM) vs. DMSO control. All images in B) were acquired and processed with identical settings. Scale bars = 5 μm.

OPP labeling of nascent polypeptides requires active protein synthesis and should be responsive to chemical inhibition of translation prior to OPP labeling. Treatment of infected HepG2 cells with pan-eukaryotic or *Plasmodium*-specific translation inhibitors recapitulated known inhibitor specificity. Acute treatment with cycloheximide, which blocks translation elongation via binding the ribosomal E-site (23) and is active against both human and *Plasmodium* translation (24, 25), results in loss of the OPP-A555 signal, indicating a dramatic drop in protein synthesis of both HepG2 and parasite (Fig. 1B). DDD107498 is a *Plasmodium*-specific translation inhibitor thought to target eEF2 (11), and treatment results in loss of OPP-A555 signal only in the parasite, with host HepG2 and parasite nascent proteome (OPP-A55 signal) clearly separable with confocal microscopy (Fig. 1B, S3). Taken together, our data suggest that OPP labeling of the nascent proteome will allow separate quantification of *Plasmodium* liver stage translation and that of the host HepG2 cells, thus opening up the study of chemical inhibitors of translation beyond the *Plasmodium* asexual blood stage.

### Quantification of the *P. berghei* and HepG2 nascent proteomes

To move from visualization to quantification of the nascent proteome, we utilized automated confocal feedback microscopy (ACFM) (26) to generate unbiased confocal image sets of single *P. berghei* liver stage parasites and the HepG2 cells immediately surrounding them (referred to as in-image HepG2). Image sets consisted of 3 separately acquired channels with anti-HSP70 marking the exoerythrocytic parasite forms (EEFs), Hoechst-labeled DNA, and OPP-A555 labeling the nascent proteome in both HepG2 and EEF. We established a CellProfiler (27) pipeline for batch image processing, in which an EEF object segmented in the anti-HSP70 image was then used to mask the other two images for further segmentation and feature extraction (Fig. S4), incuding fluorescene intensity metrics describing the magnitude of parasite translation via the OPP-A555 signal. To quantify the in-image HepG2 nascent proteome regardless of OPP-A555 signal intensity, we used segmented HepG2 nuclei from the Hoechst image to define the pixels within which to quantify OPP-A55, as this measurement is tightly correlated (R = 0.94) with the full cellular HepG2 OPP-A555 signal in control images (Fig. S5). With an image segmentation and feature extraction pipeline in place, we returned to OPP-A555 labeling controls to establish the detectable range of signal specific to the nascent proteome in both *P. berghei* EEFs and in-image HepG2 cells. Infected cells that received no OPP, but were subjected to a click labeling reaction with A555, had a larger signal than those which were OPP labeled without fluorophore conjugation. Both were extremely small, though, relative to the specific signal from parasite and HepG2 nascent proteomes, allowing us to specifically quantify translation over a range of ≥ 3 log units (Fig. S6).

### Assessing liver stage translation inhibition by compounds with diverse mechanisms of action

Having established a robust assay to quantify the *P. berghei* LS nascent proteome, we next tested a select set of compounds including 9 antimalarials and 10 pan-eukaryotic bioactive compounds for their ability to inhibit *P. berghei* LS translation. The pan-eukaryotic actives include 7 compounds that are known translation inhibitors and 3 compounds with different mechanisms of action (Table S1). While all the pan-eukaryotic compounds have demonstrated antiplasmodial activity against *P. falciparum* asexual blood stages in either growth or re-invasion assays where compounds are present throughout 48+ hours (28–34), comparable data for their liver stage activity cannot be generated due to the confounding effects that such compounds have on HepG2 cell viability (Fig. S7). To avoid confounding effects of long term treatment, we first tested compounds for ability to inhibit *P.* berghei LS translation after an acute pre-treatment of 3.5 h followed by 30 minutes of OPP labeling in the continued presence of test compound (Fig. 2A). Each of the 19 compounds were tested at micromolar concentrations expected to be saturating, but which did not induce visible HepG2 toxicity, such as cell detachment or rounding up during 4 h (Table S1). 6 of the 7 pan-eukaryotic translation inhibitors tested inhibited *P. berghei* LS translation by ≥ 90%, and, as expected, the same 6 translation inhibitors reduced HepG2 translation by ≥ 90% (Fig. 2B). In contrast, treatment with the threonyl-tRNA synthetase inhibitor borrelidin (35, 36) caused only a 53% mean reduction *P. berghei* LS translation and an 88% mean reduction in HepG2 translation (Fig. 2B). Differences in efficacy of human and *Plasmodium* translation inhibition were also detected for several other compounds. Halofuginone, an inhibitor of *P. falciparum* prolyl-tRNA synthetase (37), and emetine, which inhibits *P. falciparum* elongation (38), both displayed greater efficacy against HepG2 than *P. berghei*, while cycloheximide was slightly more effective against the parasite (Fig. 2B). Bruceantin, a translation initiation inhibitor (39), and the elongation inhibitors anisomycin and lactimidomycin (23) caused similar levels of translation inhibition between *P. berghei* and HepG2 cells (Fig. 2B).

**Figure 2.**
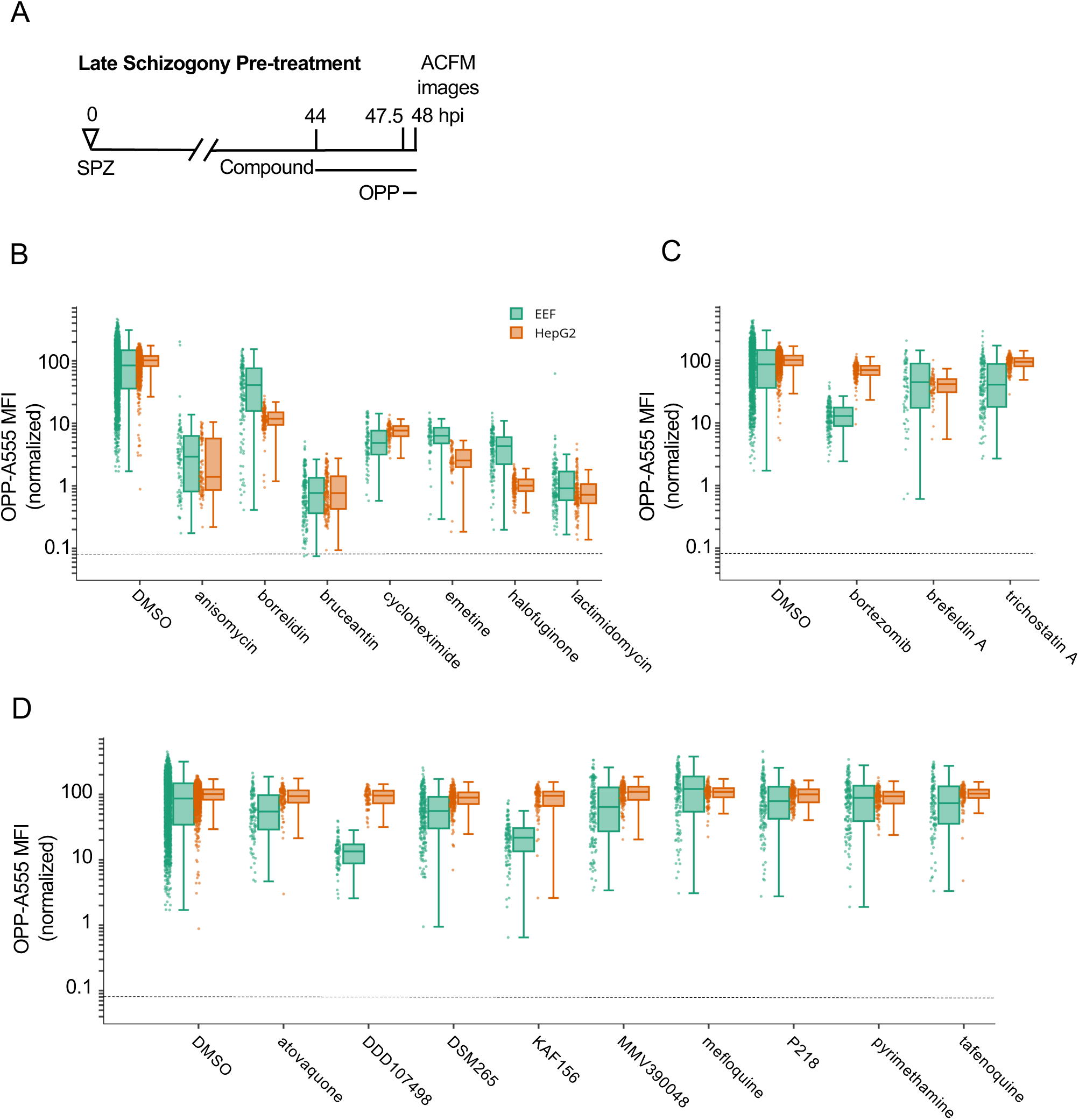
Testing select bioactive compounds for inhibition of *Plasmodium* liver stage translation. Quantification of *P. berghei* and HepG2 protein synthesis after 3.5h pre-treatment with diverse, active compounds, as schematized in A). See Table S-1 for compound details. B-D) Boxplots quantifying translation inhibition via OPP-A555 mean fluorescence intensity (OPP-A555 MFI) from all single parasite ACFM images acquired for n≥3 independent experiments, with each dot corresponding to a single EEF (green) or associated in-image HepG2 cells (orange). Specific signal cutoff (see Figure S2-3) is indicated by the dashed line. Compounds tested are known pan-eukaryotic translation inhibitors B), pan-eukaryotic inhibitors of cellular processes other than translation C), and antimalarial compounds D). Compounds were tested at 10 µM except for cycloheximide (10 µg/mL), tafenoquine (1.25 µM), mefloquine (2.5 µM), trichostatin A (5 µM) & brefeldin A (5 µg/mL).

To probe assay specificity, we tested three compounds known to be highly active against HepG2 and *Plasmodium* with cellular modes of action other than translation inhibition, which lead to complete HepG2 toxicity within 48 hours (Fig. S7). Surprisingly, the 26S proteasome inhibitor bortezomib (40) caused an 86% reduction in *P. berghei* liver stage translation, but had little effect on HepG2 translation (Fig. 2C). Trichostatin A, a histone deacetylase inhibitor (41), and Brefeldin A, which blocks the secretory pathway in *P. berghei* LS and *P. falciparum* ABS (42, 43) inhibited liver stage translation by 44% and 46% respectively (Fig. 2C and Table S1). The third group of test compounds consisted of known antimalarials, with all but mefloquine known to be active against *Plasmodium* liver stages (44). None of these antimalarials affected HepG2 translation following acute pre-treatment (Fig. 2D), but two substantially reduced liver stage translation. DDD107498 (cabamiquine, M5717), thought to act via eEF2 inhibition and a known translation inhibitor in *P. falciparum* ABS (11) inhibited liver stage translation by 86% (Fig. 2D). KAF156 (Ganaplacide), thought to affect the secretory pathway at the level of the ER or Golgi (45, 46), was unexpectedly active in the assay, reducing mean *P. berghei* translation by 76% (Fig. 2D). Three antimalarial compounds caused only slight decreases in *P. berghei* translation following a 3.5 h acute pre-treatment, including atovaquone, which targets the bc1 complex (47), DSM265, a *Plasmodium* DHODH inhibitor (48), and MMV390048, which targets *Plasmodium* PI4K (49). The remaining antimalarial compounds had little or no on parasite translation and included *Plasmodium* DHFR inhibitors pyrimethamine (50) and P218 (51), the 8-aminoquinalone tafenoquine, which lacks a clear mechanism (52), and mefloquine, thought to target *Plasmodium* blood stage feeding but also proposed to inhibit the ribosome (53, 54).

All compounds inhibiting *P. berghei* or HepG2 translation by at least 50% were considered active, and progressed to concentration-response analysis.

We initially chose to run the acute pre-treatment assay during late schizogony due to the advantages of imaging larger parasites, but found that a substantial number of control parasites had translational outputs resembling those pre-treated with translation inhibitors (Fig. 2). Given that all *Plasmodium* LSs do not successfully complete development *in vitro* (55, 56), we performed the concentration-response experiments during both early and late schizogony in parallel (Fig. 3A) to additionally probe for developmental differences in parasite translation. Using 11 paired datasets, raw mean translation intensity in 28 vs. 48 hpi parasites was significantly different while that of in-image HepG2 was not (p= 0.00019 (LS), 0.0995 (HepG2); paired t-test). We defined individual parasites as “translationally impaired” if the OPP-A555 MFI was ≤ 50% of the mean OPP-A555 of all in-plate DMSO controls, and similarly classified the in-image HepG2. Using this definition of translational impairment, there is a substantial increase in translationally impaired control LSs at 48 hpi (33.3%) vs. 28 hpi (7.7%), while a more modest shift was seen in the HepG2 (Fig. 3B). On average, parasite size was highly similar between translationally impaired and unimpaired parasites at 28 hpi, but markedly different at 48 hpi (Fig. S8 p<0.005), suggesting that translational impairment at 48 h control parasites is indicative of earlier developmental failure or growth inhibition.

**Figure 3.**
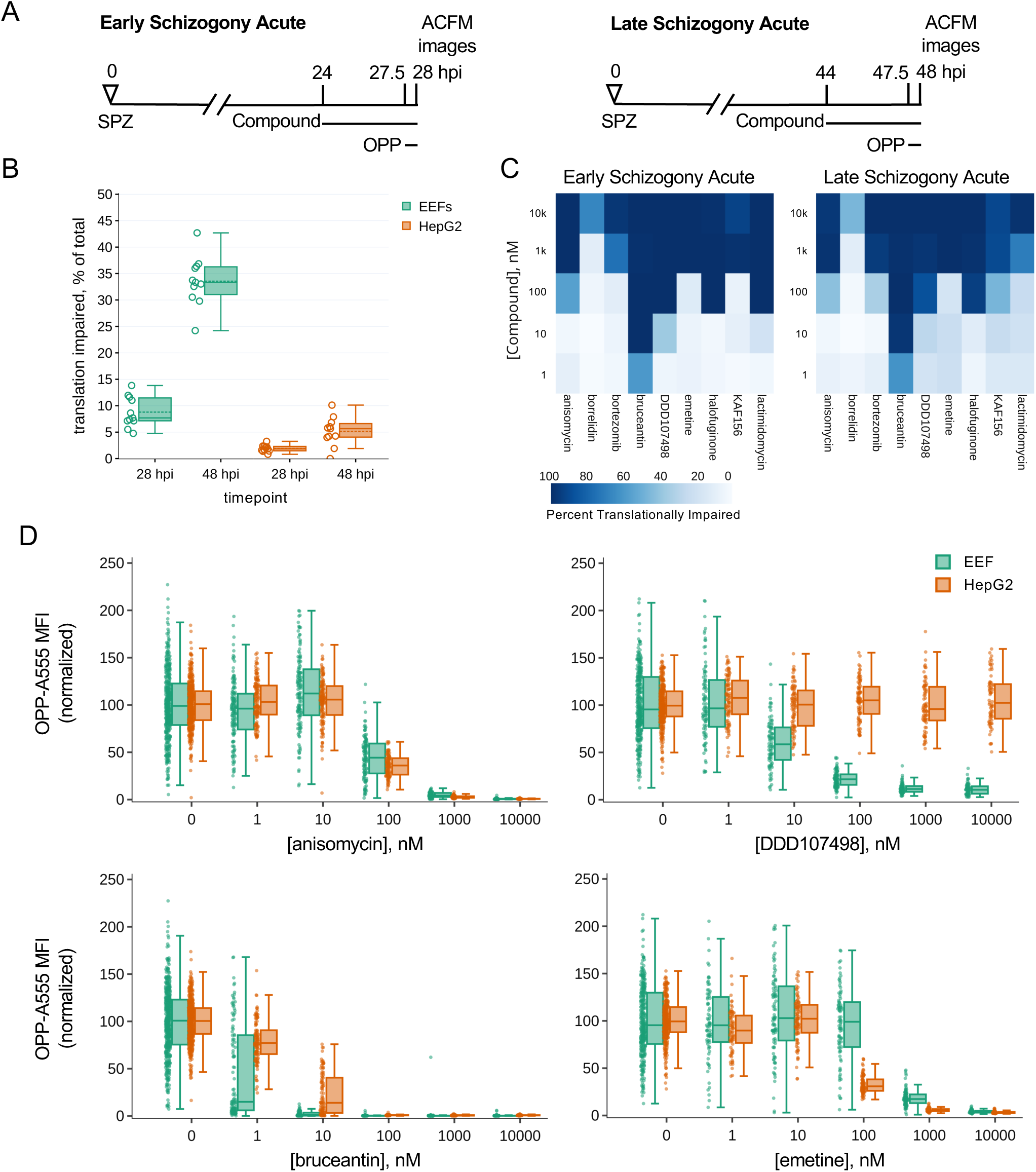
Assessing heterogeneity and potency of translation inhibition during early and late schizogony. A) Experimental schematics. B) DMSO treated control EEFs and corresponding in-image HepG2 cells were classed as translationally impaired (individual parasite OPP-A555 MFI ≤ 50% of experiment OPP-A555 mean) or unimpaired during early and late schizogony; data show the mean of 11 matched independent experiments with circles representing individual experiment values. C-D) Determining potency of translation inhibition in *P. berghei* and in-image HepG2 cells after acute pre-treatment with the inhibitors identified in Fig. 2; n≥3 independent experiments C) Percentage of single EEFs categorized as translationally impaired for inhibitors across concentrations in early and late schizogony. D) Concentration-response data in single parasites and in-image HepG2 cells for select translation inhibitors; all data normalized to mean of in-plate DMSO controls, set to 100.

Despite the marked difference in translational heterogeneity between the parasite populations at 28 and 48 hpi, both efficacy and potency of the 10 compounds active against liver stage protein synthesis were quite similar in early vs. late schizogony, reproducible across independent experiments (Figs. 3C, S9, S10). Anisomycin, which blocks elongation by occupying the A-site and preventing peptide bond formation (23, 57), has very similar potency against human and *P. berghei* translation, while DDD107498 is completely parasite-specific, as expected (Fig. 3D). Modest selectivity towards *P. berghei* is seen for bruceantin, the most potent inhibitor tested, while emetine has greater potency against HepG2 protein synthesis (Fig. 3D, S9, S10). For all pan-eukaryotic translation inhibitors tested except borrelidin, which only achieved 64% inhibition at the maximum concentration tested against early LSs, translation inhibition efficacy was similar between HepG2 and *Plasmodium* (Fig. 3D, S9, S10). Lactimidomycin lost potency against both *Plasmodium* and HepG2 translation during late schizogony (Figs. 3C, S9, S10); this likely reflects compound instability (see Methods). Anisomycin, bruceantin, cycloheximide, halofuginone, and lactimidomycin all inhibited protein synthesis in early liver stage schizonts by > 95% (Table S2) after 3.5 h of treatment, despite their varied modes/mechanisms of action. DDD107498, reached only 90.5% inhibition with the same treatment duration at the highest dose tested, despite having clearly achieved a saturating response (Fig. 3D, Table S2). Concentration-dependent inhibition of LS translation was seen for both KAF156 and bortezomib, which reached 77% and 86% inhibition, respectively (Table S2).

### Differentiating between direct and indirect inhibitors of *Plasmodium* translation

The acute pre-treatment assay was designed to maximize signal from translation inhibitors while avoiding confounding effects from HepG2 toxicity often seen with long treatment windows. However, this means that the assay should identify both direct protein synthesis inhibitors, and those that inhibit translation indirectly, e.g. compounds that induce cellular stress, leading to a signaling-based shutdown of protein synthesis via phosphorylation of eIF2α (58). To test whether our 10 active compounds are direct or indirect translation inhibitors, we ran a competition OPP assay (co-OPP), where OPP and the compound of interest are added to *P. berghei*-infected HepG2 monolayers concomitantly. Since puromycin analogues like OPP truncate a nascent polypeptide chain at the position they are incorporated, the co-OPP assay effectively means there is direct competition between the test compound and OPP to shut down translation of each nascent polypeptide at each codon (17, 20–22). Direct translation inhibitors will reduce OPP-A555 labeling of the nascent proteome competitively, while indirect translation inhibitors are expected to be inactive, or with reduced activity, in the co-OPP assay.

The co-OPP assay was first run at top concentration (see Table S1) during both early and late *P. berghei* schizogony. Strikingly, both KAF156 and bortezomib, the two unexpected actives in acute pre-treatment mode, were not competitive inhibitors of OPP labeling at either timepoint, and are thus indirect translation inhibitors (Fig. 4A-B). Here, bortezomib treatment increased translational intensity in HepG2 cell at both time points and in early *P. berghei* schizonts (Fig. 4A-B and Table S2). Anisomycin, bruceantin, cycloheximide, emetine, halofuginone, and lactimidomycin were all direct inhibitors of both *P. berghei* and HepG2 protein synthesis, while DDD107498 was a direct inhibitor of parasite translation only (Fig. 4A-B). Anisomycin and DDD107498, thought to act against the elongation step of protein synthesis, and bruceantin, which inhibits translation initiation, were selected for 5pt. 10-fold serial dilution dose response to test whether any difference in potency could be detected in co-OPP versus acute pre-treatment assays in early *P. berghei* liver stage schizonts. Bruceantin showed a clear reduction in parasite translation inhibition potency in the competition assay, with a ∼6-fold shift in EC_50_, while DDD107498 and anisomycin did not (Fig. 4C and Table S2). The success of the competition assay in identifying all known direct inhibitors of HepG2 translation, and the demonstration that these pan-eukaryotic actives are similarly direct translation inhibitors in *P. berghei* EEFs suggests that the co-OPP assay can be useful to identify unknown translation inhibitors in primary or secondary screens.

**Figure 4.**
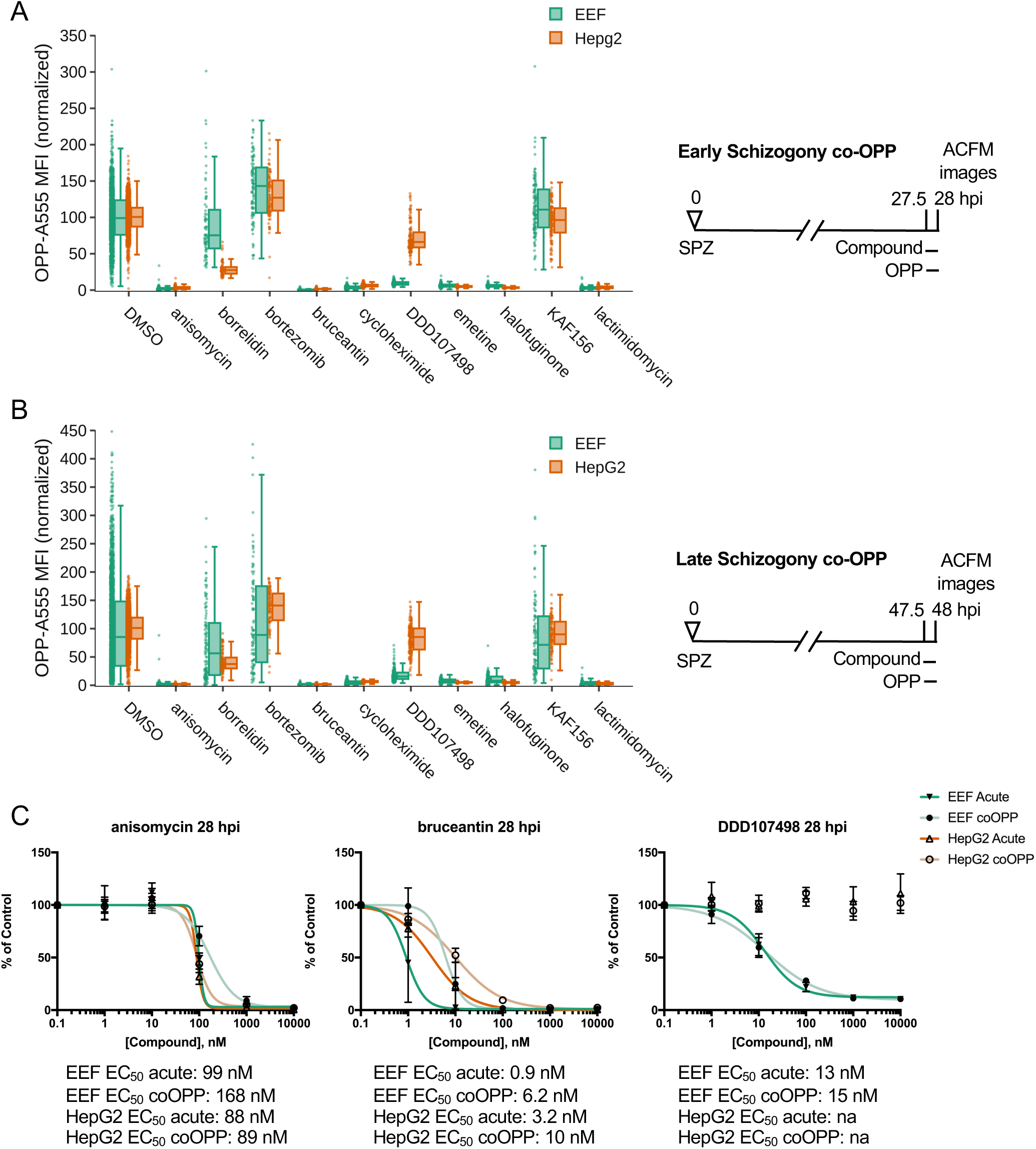
Identification of direct vs. indirect translation inhibitors. Quantification of protein synthesis in *P. berghei*-infected HepG2 cells in which active compounds at maximal concentrations were added together with OPP in early (A) and late (B) schizogony as described in figure schematics. Compound concentrations tested are the same as in Fig. 2. Each data point represents the normalized OPP-A555 mean fluorescence intensity (OPP-A555 MFI) of a single EEF or the corresponding HepG2 cells as labeled. C) Comparing coOPP and acute pre-treatment (from Fig. S3-1) concentration-response curves. All data shown was collected in n ≥ 3 independent experiments.

### Investigation of the mechanism of indirect translation inhibition by bortezomib and KAF156

Our finding that bortezomib and KAF156 similarly caused indirect translation inhibition in *P. berghei* LSs was unexpected, as they have distinct modes of action. They may, however, converge phenotypically downstream of ER stress, as bortezomib-driven accumulation of misfolded or damaged proteins in the ER causes an unfolded protein response (UPR) that is partially conserved in *Plasmodium* (59), while multiple lines of evidence indicate that KAF156 affects the parasite ER (45, 60). To investigate whether ER stress might be driving the indirect translation inhibition caused by KAF156 and bortezomib, we first investigated the phenotypic impact of both compounds on *P. berghei* LS ER structure using BiP, an HSP70 localized to the ER lumen (61), as a marker in immunofluorescence analysis (IFA). The LS schizont ER is a single, continuous structure composed of ER centers (tight accumulations of tubules) interconnected by a network of thin tubules (62) (Fig. 5A, DMSO). A 4h brefeldin A (BFA) treatment causes these centers to collapse into a single structure while the immunofluorescence intensity of anti-BiP labeling is similar to the control; DDD107498 treatment led to a similar collapse of ER centers, together with a substantial reduction in BiP IFA signal intensity localized to a single dim ER center (Fig. 5A). Bortezomib and KAF156 both altered the ER morphology profoundly, with the ER appearing to have fragmented or vesiculated throughout the EEF (Fig. 5A). Strikingly, bortezomib also caused a marked reduction in BiP signal intensity, like DDD107948, while KAF156 does not (Fig. 5A). These findings support the hypothesis that KAF156 and bortezomib could both induce ER stress leading to subsequent translational arrest in the PbLS.

**Figure 5.**
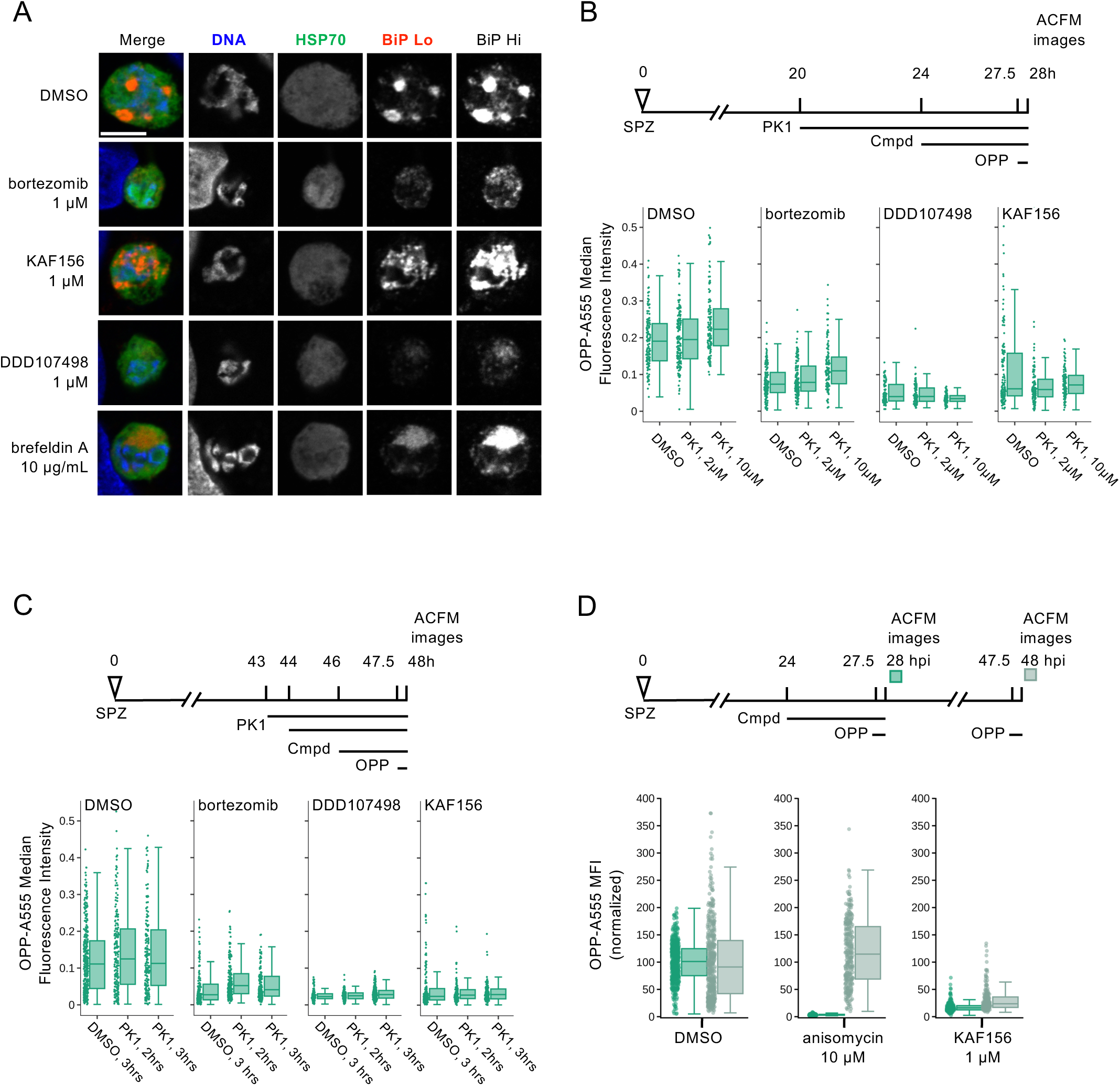
Investigating the mechanism behind indirect translation inhibition. A) Representative, single confocal images of *P. berghei* liver stage ER morphology after 4h compound treatment in early schizogony, at 28 hpi. Single channel images, all acquired with identical settings, are shown in grayscale, with merges pseudocolored as labeled; HSP70 marks the parasite and BiP specifically labels the parasite ER. Two images of BiP immunofluorescence were acquired with different gains (BiP Lo and BiP Hi) to visualize ER morphology across the range of BiP intensity observed. Scale bar = 5μm. B-D) Quantification of protein synthesis in single EEFs following treatments detailed in associated schematics. In B-C) [bortezomib] = 1 µM, [KAF156] = 0.5 µM, and [DDD107498] = 0.1 µM were used to achieve similar levels of submaximal translational inhibition in the parasites. n≥3 independent experiments. [PK1] as labelled in B), and 20 µM in C). Data in D) was normalized to the mean of the DMSO control parasites for each timepoint.

The *Plasmodium* response to ER stress appears to lack the transcriptional regulatory arm of the eukaryotic UPR (62, 63), but that which attenuates translation via eIF2α phosphorylation is present and active. Three eIF2α kinases exist in *Plasmodium* (64), with PK4 (PBANKA_1126900, PF3D7_0628200) mediating phosphorylation of eIF2α when ER stress is induced by DTT or artemisinin in *P. berghei* and *P. falciparum* asexual blood stages (65–67). *Plasmodium* PK4 appears orthologous to the human PERK kinase, and *P. falciparum* and *P. berghei* PK4 activity can be inhibited by the human PERK inhibitor GSK2606414 (PK1) (65, 66, 68). To test whether indirect translation inhibition caused by bortezomib and KAF156 was mediated by the eIF2α kinase PK4, we tested if PK1 pre-treatment could prevent translation inhibition by these compounds. DDD107498 was used as a control, since it inhibits *P. berghei* LS translation directly (Fig. 4), and PK4 inhibition should thus have no effect on its activity. We first tested 4-hour PK1 pre-treatment at 0 (DMSO control), 2 or 10 μM from 20-24 hpi, followed by the addition of KAF156 (0.5 μM), bortezomib (1 μM), DDD107498 (0.1 μM) or DMSO from 24-28 hpi, with OPP added in the final 30 minutes. These concentrations of KAF156, bortezomib and DDD107498 were chosen to induce sub-maximal translation inhibition, and showed clear, but incomplete reduction in the OPP-A555 median fluorescence intensity in single parasites (Fig. 5B). However, pre-treatment with PK1 did not prevent subsequent inhibition of EEF translation by bortezomib, KAF156, or DDD107498 (Fig. 5B). We also tested a shortened 20 μM PK1 pre-treatment and shortened KAF156 and bortezomib treatments, as prolonged PK1 treatment at this concentration leads to HepG2 cytotoxicity (not shown). Control experiments demonstrate that 2 h treatments with bortezomib or KAF156 are sufficient to induce translational arrest, but once again, PK1 was not able to prevent translation inhibition by either compound (Fig. 5C). In both PK4 inhibition protocols, in-image HepG2 translation was also quantified. Bortezomib treatment alone led to a reduction in HepG2 translation as has been previously demonstrated, and shown to be mediated by human PERK (69); PERK inhibition by PK1 pre-treatment markedly increased HepG2 translation after addition of bortezomib (Fig. S11). These results demonstrate that the indirect translation inhibition induced by KAF156 and bortezomib is not mediated by *Plasmodium* PK4. Another hypothesis for this indirect translation inhibition is that it reflects a rapid parasite death process. If so, the translation inhibition should not be reversible. We tested this directly by comparing reversibility of the translation inhibition induced by 4 h KAF156 and anisomycin treatments in early schizogony. Anisomycin-induced translation inhibition is reversible in human cells (57), and the ∼95% inhibition of *P. berghei* liver stage translation was completely reverted 20h after compound washout (Fig. 5D). KAF156 treatment induced weaker translation inhibition (∼85%) compared to anisomycin but showed very little recovery of translation 20 h after washout. The irreversibility of the translation inhibition after washout suggests that KAF156 treatment causes rapid parasite death.

### Testing uncharacterized *P. berghei* liver stage active compounds for the ability to inhibit protein synthesis

Finally, to investigate the utility of this assay for identifying novel *Plasmodium* protein synthesis inhibitors, we tested 6 compounds from the MMV Malaria Box that are active against *P. berghei* liver stages and phenotypically similar to DDD107498 in 48h luciferase assays (70). Acute pre-treatment with MMV019266 reduced EEF translation by 87% (Fig. 6A). The remaining compounds were much less active, with MMV665940, MMV007116, and MMV006820 causing roughly 30% reduction in PbLS translation, MMV006188 causing a 19% reduction, and MMV011438 having no effect (Fig. 6A). MMV019266 similarly inhibited LS translation at both 1 and 10 μM during early and late schizogony (Figs. 6B-C, S12, Table S2). MMV019266 had EC_50_ values of 373 and 289 nM at 28- and 48 hpi, respectively in the acute pre-treatment assay (Fig. S12 and Table S2). MMV019266 was also capable of inhibiting *P. falciparum* ABS translation in intact schizonts, with the degree of inhibition similar to that seen with 10 μM DDD107498, but greater than 20 nM DDD107498 and less than 100 nM bruceantin (Fig. S13). The co-OPP assay demonstrated that MMV019266 is a direct protein synthesis inhibitor, causing 77% and 72% reductions in PbLS translational intensity during early and late schizogony respectively (Fig. 6D and Table S2). Identification of MMV019266 as a direct translation inhibitor in both blood stage and liver stage parasites highlights the utility of the *P. berghei* LS OPP assay to antimalarial drug discovery.

**Figure 6.**
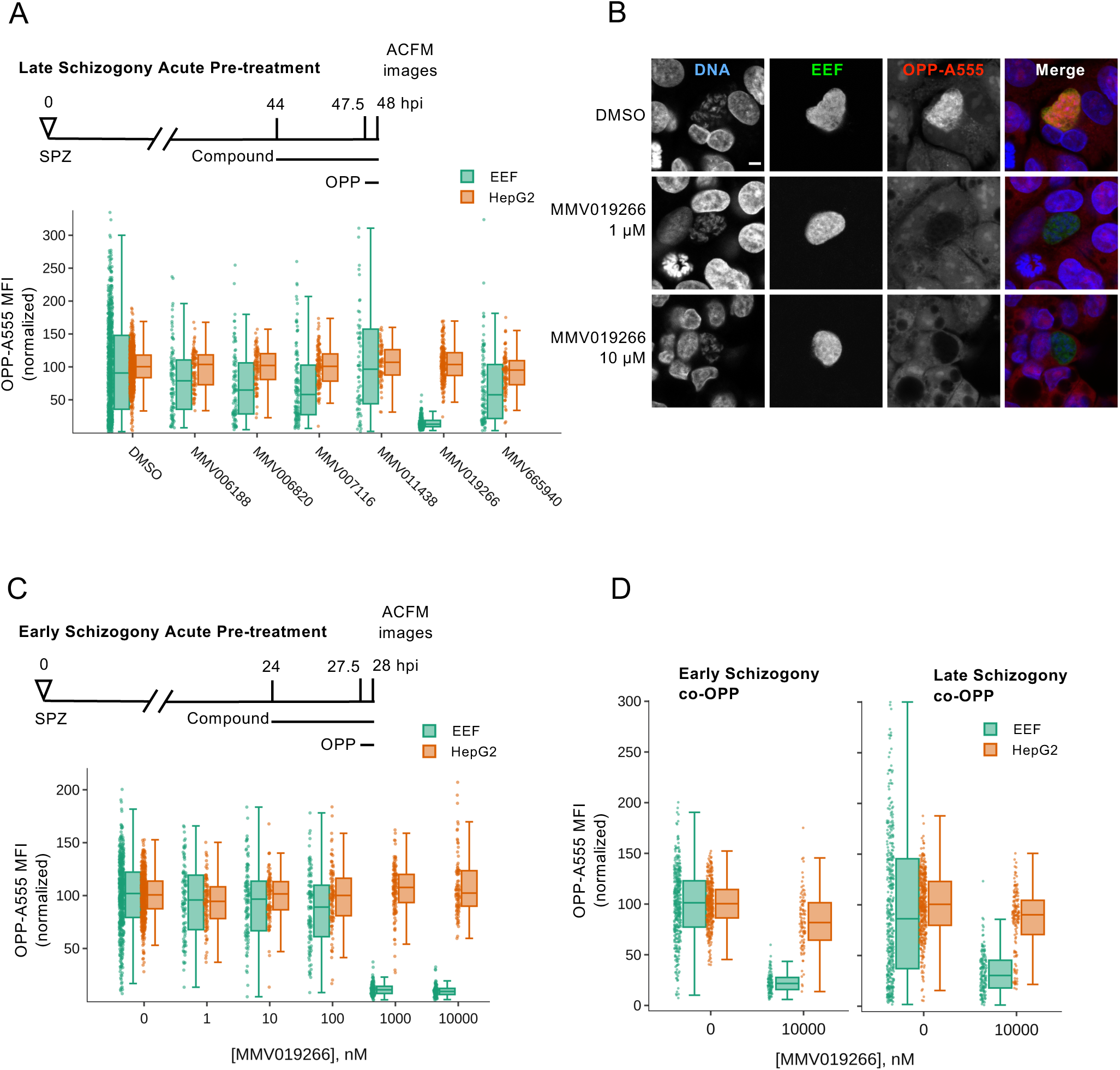
Characterization of MMV019266 inhibition of *P. berghei* liver stage translation. A) Select liver stage actives from the Malaria Box were tested at 10 µM for ability to inhibit *P. berghei* liver stage translation following acute pre-treatment in late schizogony. B) Representative single confocal images of OPP-A55 labeling after 4h acute pre-treatments with MMV019266 vs. control in late schizogony; merges are pseudo colored as indicated with parasite (EEF) immunolabeled with α-HSP70, and DNA stained with Hoechst. Scale bar = 5 μm. C-D) Quantification of protein synthesis in *P. berghei-*infected HepG2 cells; compound treatments as described. Each data point represents OPP-A555 mean fluorescence intensity (MFI) normalized to in-plate controls; n≥3 independent experiments.

## Discussion

Our results demonstrate the feasibility and utility of single cell image-based quantification of protein synthesis in an intracellular parasite that resides in a translationally active host cell, and open up the study of *Plasmodium* liver stage translation for drug discovery applications and in the native developmental context. To date, studies of the mode of action and target of antimalarial compounds largely rely on studies in *P. falciparum* asexual blood stages (ABS), with the assumption that it will be the same in other stages and species in which a compound has antiplasmodial activity. Molecular targets of antimalarials compounds have been identified and validated in *P. falciparum* ABS through *in vitro* evolution of drug resistance, cellular thermal shift assays, chemoproteomics, metabolic profiling, and a variety of reverse genetics approaches to produce modified parasite lines (71–74). Re-use of these same evolved or genetically modified parasite lines, metabolic profiling and assays quantifying intracellular ionic concentrations all support further understanding of antimalarial compound modes of action in ABS (75–77). Protein synthesis in ABS has long been quantified by feeding with radiolabelled amino acids and more recently *P. falciparum* ABS lysate assays detecting the translation of a single model transcript encoding a luciferase enzyme have been used to screen several small compound libraries and characterize the activity of known pan-eukaryotic translation inhibitors (14, 15, 78).

An overarching difficulty in quantification of conserved biochemical or cellular processes like protein synthesis in *Plasmodium* liver stages is the dominant contribution of hepatocytes to any signal from an infected monolayer. In ABS, this problem is easily overcome, as saponin lysis of infected RBC cultures has long been recognized to allow parasite purification (79), and mature human erythrocytes lack most core cellular processes, e.g. protein synthesis, allowing *Plasmodium* translation to be quantified directly in bulk ABS cultures or lysates. Inability to physically isolate liver stage parasites or isolate the parasite signal from that of the hepatocytes prevents the use of these approaches in *Plasmodium* liver stages currently. Here, we overcome this limitation by using computational separation of the combined fluorescent signal of the nascent proteome of infected HepG2 monolayers into separate *P. berghei* and hepatoma cells signals in ACFM-acquired image sets. The specificity of this approach is clear from *Plasmodium*-specific inhibition of translation by DDD107498, and our ability to detect differential inhibition of *H. sapiens* vs. *P. berghei* translation with pan-eukaryotic inhibitors like emetine and bruceantin means that both host and parasite nascent proteomes can be quantified in parallel, allowing determination of a compound’s LS translation inhibition efficacy and selectivity in a single well. Similar quantitative bioimaging strategies may prove useful for drug discovery efforts with other eukaryotic parasites residing in translational active host cells.

One attractive feature of targeting *Plasmodium* translation is that such inhibitors would be predicted to have multistage activity, as has been demonstrated for DDD107498 and a variety of tRNA synthetase inhibitors ((11, 80). Though our data show that DDD107498 LS translation inhibition potency is 13-15 nM, clearly less than its ∼1-2 nM LS antiplasmodial potency in standard 48 h LS biomass assays, it is clearly a concentration-dependent translation inhibitor. It is striking, though, that at 1 nM we detect no clear translation inhibition at all, and the slope is very shallow, with saturating effects only seen at 1000 nM, and the percent max translation inhibition is clearly less than for other parasite-active compounds. These effects seem unlikely to be time dependent, as we show nearly identical translational responses to DDD107498 in acute pre-treatment and co-OPP assays. Incomplete translation inhibition with saturating doses of DDD107498 was also seen in *P. falciparum* ABS (11), and it will be a future challenge to determine how much translation inhibition is required for DDD107498 antiplasmodial activity. Consistent with the hypothesis that translation inhibitors should be multistage actives, we show concentration dependent LS inhibition for the elongation inhibitors anisomycin, lactimidomycin, emetine, and cycloheximide, the initiation inhibitor bruceantin, and the tRNA synthetase inhibitor halofuginone. Only borrelidin, a known inhibitor of threonyl-tRNA synthetase (ThrS) in both prokaryotes and eukaryotes (35), failed to show concentration dependent translation inhibition activity, and was only partially effective in our acute pre-treatment assay at the highest concentration tested (10 μM), which is at odds with the low nanomolar antiplasmodial potency of borrelidin (29, 36, 81–83), where reported IC50s range from 0.07 nM to 1.9 nM. We tested borreldin in a 48h LS live luciferase assay, but all concentrations that reduced parasite biomass also showed effects on the HepG2 monolayer (data not shown), so it is unclear if borrelidin has any direct antiplasmodial activity against the *P. berghei* LS. Species specific differences in activity should not be the cause, as borrelidin was active against ABS of both human and murine *Plasmodium* spp., and was an effective antimalarial in murine infection models (29, 36, 81, 82). Furthermore, this disconnect is not easily explained by stage specific differences in the target enzyme expression or activity, as *Plasmodium* parasites encode only a single copy of ThrS (84), which is likely required for protein synthesis in both the cytoplasm and apicoplast (58, 85). Enzymatic evidence clearly shows that borrelidin is active against recombinant PfThrS *in vitro* (36), but the cellular evidence in support of borrelidin targeting *Plasmodium* ThrS was a modest shift in the *P. falciparum* ABS growth inhibition EC_50_ when an excess of exogenous free L-threonine in growth media (81). Evolved *in vitro* resistance to borrelidin has not been reported to date. Given that compound efficacy against recombinant protein is not always a reliable indicator of *in vivo* antimalarial mechanism, as with triclosan (86), it will be important to clarify that ThrS is indeed the relevant antimalarial target of borrelidin, and if so, understand why it is not effective against *P. berghei* liver stages *in vitro*.

The image-based OPP assay appears to have some advantages relative to the lysate assay, PfIVT, (14, 15, 78) in testing antiplasmodial compounds of unknown mechanism for translation inhibition activity. Given the liquid handling requirements for the OPP assay, it is ideally suited for use with adherent cells, and thus liver stage parasites, and while our current 96wp format is sufficient for testing of compounds of interest as we demonstrate here, we are miniaturizing the assay to 384wp format for medium throughput use. Image-based assays have the advantage of inherent ability to investigate ground-truth of translation inhibition metrics obtained via segmentation and feature extraction, as metadata links the original, unaltered image set to extracted features (87), while a lysate-based assay lacks inherent ground truth, and may require a secondary counterscreen to triage compounds against the translated reporter enzyme, as for PfIVT (78). Translation is a complex process, requiring spatial coordination of hundreds of gene products (58, 88) to produce new proteins from thousands of mRNAs, and the image-based OPP assay captures changes in output of the entire, native nascent proteome. A lysate-based assay using translation of a single exogenous mRNA as a readout reduces this complexity substantially, and may fail to identify compounds that active translation inhibitors *in cellulo*. Perhaps this occurred with MMV019266, which was not identified as an active compound in the PfIVT screen of the Malaria Box (14). We tested 6 compounds identified from the MMV Malaria Box as LS active with a 48h biomass assay phenotype indicative of early liver stage arrest (70), the same as for DDD107498, which we could source commercially. While 5 were inactive, the thienopyrimidine MMV019266 was identified as a direct translation inhibitor in *P. berghei* LS, and we demonstrated that it also inhibits *P. falciparum* translation in blood stage schizonts. MMV019266 is known to have antiplasmodial activity against a variety of species and life cycle stages, including *P. vivax* schizonts and *P. falciparum* gametocytes (70, 89–93), and was predicted to target hemoglobin catabolism based on metabolic fingerprinting (94). During the preparation of this manuscript, three related thienopyrimidines were reported to target the *P. falciparum* cytoplasmic isoleucyl tRNA synthetase (PfcIRS) based on mutations evolved *in vitro* in resistant lines and confirmed in conditional PfcIRS knockdowns and gene-edited parasite lines (95). These results highlight the value of the *P. berghei* liver stage OPP assay for identification of multistage *Plasmodium* translation inhibitors.

While our focus here has been using the quantitative image-based OPP assay to identify novel *Plasmodium* translation inhibitors and validate this mode of action in the liver stage for known inhibitors of *P. falciparum* blood stage translation, the flexibility of the assay and power of single cell data suggest it may prove quite useful in key applications beyond this. While our quantitative work utilized LS schizonts, we demonstrated specific nascent proteome signal in sporozoites through to monolayer merozoites, and intriguing changes in the subcellular localization of the nascent proteome seem to occur during *P. berghei* LS development. The difference in the fraction of translationally impaired parasites in 28 vs. 48 hpi LS schizonts suggests that translational intensity may vary during liver stage development in populations, as it clearly does in individuals at both timepoints. Our identification of Kaf156 and bortezomib as indirect translation inhibitors not under the control of the eIF2α kinase PK4, but likely causing rapid killing of *P. berghei* LS schizonts, indicates the potential of using translational output as a biomarker for liver stage parasite viability, something that remains lacking in the liver stage toolkit (96). The labeling protocol adapts easily to *P. falciparum* blood stages, as we demonstrate, though throughput is limited by the non-adherent erythrocytes, which also complicates high content quantitative imaging. While the throughput problem will be challenging to solve for medium to high throughput drug discovery, flow cytometry may be better suited to quantification of translation inhibition in based on fluorescent labeling of the *P. falciparum* nascent proteome, as has recently been done to characterize novel tyrosine-RNA synthetase inhibitors (97). As OPP labeling of the nascent proteome requires no transgenic technology, it should be readily adaptable to critical drug discovery challenges such as testing target engagement and potency of translation inhibitors with diverse molecular mechanisms of action against field isolates of *P. falciparum* and *P. vivax*. Our quantitative bioimaging workflow should be repurposable for interrogation of translation inhibitors in *P. falciparum* and *P. vivax* liver stages and capable of integration into existing image based-screening platforms (98), and may have particular value in examining the role translation plays in formation of dormant hypnozoites and their reactivation.

## Materials and Methods

### HepG2 culture and *P. berghei* sporozoite isolation and infection

HepG2 human hepatoma cells were cultured in Dulbeco’s Modified Eagle Medium (DMEM) (Gibco 10313-021) supplemented with 10% (v/v) FBS, 1% (v/v) GlutaMAX (Gibco 35050-061), 1% (v/v) Penicillin-Streptomycin (Gibco 15140-122) and maintained at 37°C, 5% CO_2_. *Plasmodium berghei* sporozoites, expressing firefly luciferase-GFP fusion protein under the control of the exoerythrocytic form 1a (EEF1a) promoter (99), were isolated from the salivary glands of infected *Anopheles stephensii* mosquitos (NYU and UGA insectaries). Sporozoites were counted and diluted into infection DMEM (iDMEM) – cDMEM further supplemented with 1% (v/v) Penicillin-Streptomycin-Neomycin (Gibco 15640-055), 0.835µg/mL Amphotericin B (Gibco 15290-018), 500µg/mL kanamycin (Corning 30-006-CF), and 50µg/mL gentamycin (Gibco 15750-060), added to HepG2 monolayers, centrifuged at 3000 rpm for 5 minutes, and incubated in cell culture conditions for 2 hours before PBS washing and iDMEM replenishment for infections proceeding on glass coverslips. For infections in 96 well plates (Greiner 655098), infected HepG2 monolayers were detached at 2 hpi using TrypLE Express (Gibco 12605-028), washed, counted, and re-seeded into 96 well plates.

### Compound handling and treatment

Compound stocks prepared from powder were solubilized in DMSO (Sigma-Aldrich D2650), aliquoted, and stored at −20°C. For acute pre-treatments, infected cells were treated for 3.5 h prior to 30-minute OPP labeling in continued presence of compound. For coOPP assays, compound and OPP were applied simultaneously for 30 minutes. Concentration-response experiments were performed with 5 points in 10-fold serial dilutions, and equimolar DMSO concentrations (0.001% v/v) were maintained across all treatments and controls. Compounds prepared in iDMEM were stored at 4°C and used within 24 h, e.g. a single dilution series was prepared and used for both the 24- & 44- hpi additions.

### OPP labeling and fluorophore addition

A 20 mM stock of O-propargyl puromycin (OPP) (Invitrogen C10459) in DMSO was diluted to label cells with 20 µM OPP for 30 mins at 37°C, according to the manufacturer’s recommended protocol, before 15-minute fixation with PFA (Alfa Aesar 30525-89-4) diluted to 4% in PBS. Copper-(I)-catalyzed cycloaddition of Alexafluor555 picolyl azide to OPP-labeled polypeptides was performed using Invitrogen Click-iT Plus AF555 (Invitrogen C10642) according to the manufacturer’s recommendations, with a 1:4 Cu_2_SO_4_ to copper protectant ratio. 27µL of reaction mix was added to 96-wp wells, with 25 µL used for each glass coverslip, inverted on parafilm.

### Immunofluorescence

EEFs were immunolabeled using anti-PbHSP70 (2E6 mouse mAb) (100) (1:200), followed by donkey anti-mouse Alexafluor488 (Invitrogen A21202). In Fig. 1A, goat anti-UIS4 (Sicgen AB0042-500) (1:1000) was used to mark the newly invaded sporozoites, followed by donkey anti-goat 488 (Invitrogen A32814). To visualize the parasite ER in Fig. 5A, rabbit polyclonal anti-BiP (1:600, GenScript) serum, raised against the C terminal polypeptide CGANTPPPGDEDVDS from PBANKA_081890 was used with donkey anti-rabbit Alexafluorr555 (Invitrogen A31572) as the secondary. DNA was stained with Hoescht 33342 (Thermo Scientific 62249) (1:1000). Antibodies were prepared in 2% BSA in PBS, with secondary antibodies used at a 1:500 dilution.

### Plasmodium falciparum culture

*P. falciparum* 3D7 parasites were cultured as previously described (101). In short, parasites were cultured in human AB^+^ erythrocytes (Interstate Blood Bank, Memphis, TN, USA) at 3 – 10% parasitemia in complete culture medium (5% hematocrit). Complete culture medium consisted of RPMI 1640 medium (Gibco #32404014) supplemented with gentamicin (45 µg/ml final concentration; Gibco #15710064), HEPES (40 mM; Fisher #BP3101), NaHCO_3_ (1.9 mg/ml; Sigma #SX03201), NaOH (2.7 mM; Fisher #SS266-1), hypoxanthine (17 µg/ml; Alfa Aesar #A11481-06), L-glutamine (2.1 mM; Corning #25005Cl), D-glucose (2.1 mg/ml; Fisher #D16-1), and 10% (vol/vol) human AB^+^ serum (Valley Biomedical #HP1022). Parasites were cultured at 37°C in an atmosphere of 5% O_2_, 5% CO_2_, and 90% N_2_.

### *Plasmodium falciparum* blood stage immunofluorescence and OPP-A555 labeling

*P. falciparum*-infected erythrocytes (iRBCs) in mixed culture were labeled using 20 µM OPP (Invitrogen C10459) at 37°C for 30 minutes, pelleted and washed with PBS before being resuspended in 1 mL of 4% PFA (Electron Microscopy Sciences 30525-89-4) + 0.0075% glutaraldehyde (Sigma G6257) in PBS for 30 minutes at RT. Fixed iRBCs were pelleted and washed twice with PBS prior to permeabilization in 0.1% Triton X-100 (9002-93-1) in PBS for 10 minutes. Permeabilized iRBCs were washed twice in PBS, click-labeled as described for infected HepG2 monolayers, then pelleted, washed once with PBS and Hoechst-labelled for 30 minutes at RT. iRBCs were then pelleted, washed, and re-suspended in PBS before imaging.

### HepG2 viability assay

Non-infected HepG2 cells were treated with 10-point, 3-fold, serial dilutions with maximal concentrations of 10µM, except for cycloheximide (10µg/mL), GSK260414 (50µM), and emetine (25µM). At 46 hours post-treatment, AlamarBlue cell viability reagent (Invitrogen A50100) was applied at a 1X final concentration and incubated for one hour prior to measuring fluorescence at 590nm using a microplate reader (CLARIOstar, BMG LABTECH).

### Image acquisition

Images were acquired on a Leica SP8 confocal microscope using an HC PL APO 63x/1.40 oil objective for glass coverslips and an HC PL APO 63x/1.40 water objective for 96-well μclear plates. Images in Figures 1A, 5C, 7B, S1, S2, and S3 were acquired manually and processed using ImageJ (102). All other images, and all used for quantitative analysis, were acquired using automated confocal feedback microscopy (ACFM) (26). Briefly, MatrixScreener is used to define a patterned matrix for acquisition of non-overlapping, low-resolution images of the *P. berghei-*infected HepG2 monolayer. After each image is acquired, online image segmentation and ID of parasites, defined by PbHSP70 signal, is performed utilizing custom modules (https://github.com/VolkerH/MatrixScreenerCellprofiler/wiki) integrated into a CellProfiler version 2.0.11710 pipeline (27). The x-y coordinates of each parasite found are then used by MatrixScreener to sequentially image each individual parasite in high resolution, with an automated z-stack maximizing PbHSP70 intensity to identify the z coordinate, followed by sequential acquisition of Hoechst, PbHSP70, and OPP-A55 images. This process iterates until all parasites in the predefined matrix of the infected monolayer have been imaged.

### Image segmentation, feature extraction, and data cleaning

Batch image segmentation and feature extraction were performed in Cell Profiler (v2.1.1 rev6c2d896) (27); see Fig. S4 for the workflow. Briefly, EEF objects were identified using a global Otsu segmentation of the PbHSP70 image. The EEF object was shrunk by two pixels to ensure exclusion of HepG2-associated signal, and used to mask the OPP-A555 image to quantify *P. berghei* translation via OPP-A555 fluorescence intensity features. Conversely, the EEF object was expanded by two pixels and used as an inverse mask for the Hoechst image to segment HepG2 nuclei. All in-image HepG2 nuclei were unified into a single object, then its OPP-A555 fluorescence intensity features were used to quantify HepG2 translation. All features extracted were then analyzed using KNIME (103). ACFM image sets were computationally cleaned of image that did not contain a single true EEF in a HepG2 monolayer by removing data from those in which: more than one EEF object was identified, the EEF object identified did not contain a DNA signal, and no HepG2 nuclei were identified. EEF object form factor was used to identify rare instances of segmentation failure in which two parasites were segmented as a single EEF object; images sets corresponding to form factor outliers (> 1.5x IQR) were visually inspected and removed if they did not contain a single, true EEF. Finally, focus score features for both the PbHSP70 and DNA images were used to exclude any image set where focus score <1.5 IQR. Data cleaning was carried out per experiment, and resulted, on average, in the removal of 1.45% of the total data.

### Concentration response curve fitting and statistics

Concentration-response analysis was performed using 4 parameter non-linear regression curve fitting in GraphPad Prism (Version 7.0d), with the top of the curve fixed at 100, and −10< hill slope< 0. When maximal effect was reached with ≥2 concentrations tested, the bottom of the curve was fit open; if no such plateau was achieved, the curve was fit with maximal effect constrained to 0. EC_50_ and 95% CI were determined for each compound from ≥ 3 independent experiments. All other data and statistical analyses were performed in KNIME.

## Data availability

Image datasets are available upon request to the corresponding author.

## Acknowledgements

This work was supported by National Institutes of Health grant R21AI149275 to KKH. JLM was supported by a South Texas Center for Emerging Infectious Diseases fellowship. ABR was supported by Graduate Research in Immunology Program training grant NIH T32 AI138944. We thank the New York University Insectary and the University of Georgia SporoCore for providing *P. berghei*-infected mosquitos. KKH conceived the project and designed experiments. JLM, WS, ABR, and KKH performed experiments. JLM and KKH analyzed the data. EMB and KKH supervised the work in their respective laboratories. JLM and KKH drafted the manuscript. All authors participated in review and editing of the manuscript, and approved the submitted version.

## Notes

### Competing Interest Statement

The authors have declared no competing interest.

## Works Cited

1. World Health Organizaiton W. 2021. World malaria report 2021. World Health Organization, Geneva.

2. Blasco B, Leroy D, Fidock DA. 2017. Antimalarial drug resistance: linking Plasmodium falciparum parasite biology to the clinic. Nature Medicine 23:917–928.

3. Witkowski B, Khim N, Chim P, Kim S, Ke S, Kloeung N, Chy S, Duong S, Leang R, Ringwald P, Dondorp AM, Tripura R, Benoit-Vical F, Berry A, Gorgette O, Ariey F, Barale J-C, Mercereau-Puijalon O, Menard D. 2013. Reduced artemisinin susceptibility of Plasmodium falciparum ring stages in western Cambodia. Antimicrobial agents and chemotherapy 57:914–923.

4. Uwimana A, Legrand E, Stokes BH, Ndikumana J-LM, Warsame M, Umulisa N, Ngamije D, Munyaneza T, Mazarati J-B, Munguti K, Campagne P, Criscuolo A, Ariey F, Murindahabi M, Ringwald P, Fidock DA, Mbituyumuremyi A, Menard D. 2020. Emergence and clonal expansion of in vitro artemisinin-resistant Plasmodium falciparum kelch13 R561H mutant parasites in Rwanda. Nature Medicine 26:1602–1608.

5. Phillips MA, Burrows JN, Manyando C, van Huijsduijnen RH, Van Voorhis WC, Wells TNC. 2017. Malaria. Nat Rev Dis Primers 3:17050.

6. Burrows JN, Burlot E, Campo B, Cherbuin S, Jeanneret S, Leroy D, Spangenberg T, Waterson D, Wells TN, Willis P. 2014. Antimalarial drug discovery - the path towards eradication. Parasitology 141:128–139.

7. Venugopal K, Hentzschel F, Valkiūnas G, Marti M. 2020. Plasmodium asexual growth and sexual development in the haematopoietic niche of the host. Nature Reviews Microbiology 18:177–189.

8. Prudêncio M, Rodriguez A, Mota MM. 2006. The silent path to thousands of merozoites: the Plasmodium liver stage. Nature Reviews Microbiology 4:849–856.

9. Matz JM, Beck JR, Blackman MJ. 2020. The parasitophorous vacuole of the blood-stage malaria parasite. Nature Reviews Microbiology 18:379–391.

10. Wilson DN. 2009. The A–Z of bacterial translation inhibitors. Critical Reviews in Biochemistry and Molecular Biology 44:393–433.

11. Baragana B, Hallyburton I, Lee MC, Norcross NR, Grimaldi R, Otto TD, Proto WR, Blagborough AM, Meister S, Wirjanata G, Ruecker A, Upton LM, Abraham TS, Almeida MJ, Pradhan A, Porzelle A, Luksch T, Martinez MS, Luksch T, Bolscher JM, Woodland A, Norval S, Zuccotto F, Thomas J, Simeons F, Stojanovski L, Osuna-Cabello M, Brock PM, Churcher TS, Sala KA, Zakutansky SE, Jimenez-Diaz MB, Sanz LM, Riley J, Basak R, Campbell M, Avery VM, Sauerwein RW, Dechering KJ, Noviyanti R, Campo B, Frearson JA, Angulo-Barturen I, Ferrer-Bazaga S, Gamo FJ, Wyatt PG, Leroy D, Siegl P, Delves MJ, Kyle DE, et al. 2015. A novel multiple-stage antimalarial agent that inhibits protein synthesis. Nature 522:315–20.

12. Pham JS, Dawson KL, Jackson KE, Lim EE, Pasaje CF, Turner KE, Ralph SA. 2014. Aminoacyl-tRNA synthetases as drug targets in eukaryotic parasites. Int J Parasitol Drugs Drug Resist 4:1–13.

13. Chhibber-Goel J, Yogavel M, Sharma A. 2021. Structural analyses of the malaria parasite aminoacyl-tRNA synthetases provide new avenues for antimalarial drug discovery. Protein Science 30:1793–1803.

14. Ahyong V, Sheridan CM, Leon KE, Witchley JN, Diep J, DeRisi JL. 2016. Identification of Plasmodium falciparum specific translation inhibitors from the MMV Malaria Box using a high throughput in vitro translation screen. Malar J 15:173.

15. Sheridan CM, Garcia VE, Ahyong V, DeRisi JL. 2018. The Plasmodium falciparum cytoplasmic translation apparatus: a promising therapeutic target not yet exploited by clinically approved anti-malarials. Malaria Journal 17:465.

16. Forte B, Ottilie S, Plater A, Campo B, Dechering KJ, Gamo FJ, Goldberg DE, Istvan ES, Lee M, Lukens AK, McNamara CW, Niles JC, Okombo J, Pasaje CFA, Siegel MG, Wirth D, Wyllie S, Fidock DA, Baragaña B, Winzeler EA, Gilbert IH. 2021. Prioritization of Molecular Targets for Antimalarial Drug Discovery. ACS Infectious Diseases 7:2764–2776.

17. Azzam ME, Algranati ID. 1973. Mechanism of puromycin action: fate of ribosomes after release of nascent protein chains from polysomes. Proceedings of the National Academy of Sciences of the United States of America 70:3866–3869.

18. Barrett RM, Liu H-w, Jin H, Goodman RH, Cohen MS. 2016. Cell-specific Profiling of Nascent Proteomes Using Orthogonal Enzyme-mediated Puromycin Incorporation. ACS Chemical Biology 11:1532–1536.

19. David A, Dolan BP, Hickman HD, Knowlton JJ, Clavarino G, Pierre P, Bennink JR, Yewdell JW. 2012. Nuclear translation visualized by ribosome-bound nascent chain puromycylation. The Journal of cell biology 197:45–57.

20. Nathans D. 1964. PUROMYCIN INHIBITION OF PROTEIN SYNTHESIS: INCORPORATION OF PUROMYCIN INTO PEPTIDE CHAINS. Proceedings of the National Academy of Sciences of the United States of America 51:585–592.

21. Yarmolinsky MB, Haba GL. 1959. INHIBITION BY PUROMYCIN OF AMINO ACID INCORPORATION INTO PROTEIN. Proceedings of the National Academy of Sciences of the United States of America 45:1721–1729.

22. Liu J, Xu Y, Stoleru D, Salic A. 2012. Imaging protein synthesis in cells and tissues with an alkyne analog of puromycin. Proc Natl Acad Sci U S A 109:413–8.

23. Garreau de Loubresse N, Prokhorova I, Holtkamp W, Rodnina MV, Yusupova G, Yusupov M. 2014. Structural basis for the inhibition of the eukaryotic ribosome. Nature 513:517–522.

24. Schneider-Poetsch T, Ju J, Eyler DE, Dang Y, Bhat S, Merrick WC, Green R, Shen B, Liu JO. 2010. Inhibition of eukaryotic translation elongation by cycloheximide and lactimidomycin. Nat Chem Biol 6:209–217.

25. Schnell JV, Siddiqui WA. 1972. The Effects of Antibiotics on 14C-isoleucine Incorporation by Monkey Erythrocytes Infected with Malarial Parasites. Proceedings of the Helminthological Society of Washington 39:201–203.

26. Tischer C, Hilsenstein V, Hanson K, Pepperkok R. 2014. Chapter 26 - Adaptive fluorescence microscopy by online feedback image analysis, p 489-503. In Waters JC, Wittman T (ed), Methods in Cell Biology, vol 123. Academic Press.

27. Carpenter AE, Jones TR, Lamprecht MR, Clarke C, Kang IH, Friman O, Guertin DA, Chang JH, Lindquist RA, Moffat J, Golland P, Sabatini DM. 2006. CellProfiler: image analysis software for identifying and quantifying cell phenotypes. Genome Biol 7:R100.

28. Ekong RM, Kirby GC, Patel G, Phillipson JD, Warhurst DC. 1990. Comparison of the in vitro activities of quassinoids with activity against Plasmodium falciparum, anisomycin and some other inhibitors of eukaryotic protein synthesis. Biochemical pharmacology 40:297–301.

29. Otoguro K, Ui H, Ishiyama A, Kobayashi M, Togashi H, Takahashi Y, Masuma R, Tanaka H, Tomoda H, Yamada H, Omura S. 2003. In vitro and in vivo antimalarial activities of a non-glycosidic 18-membered macrolide antibiotic, borrelidin, against drug-resistant strains of Plasmodia. J Antibiot (Tokyo) 56:727–9.

30. Keller TL, Zocco D, Sundrud MS, Hendrick M, Edenius M, Yum J, Kim Y-J, Lee H-K, Cortese JF, Wirth DF, Dignam JD, Rao A, Yeo C-Y, Mazitschek R, Whitman M. 2012. Halofuginone and other febrifugine derivatives inhibit prolyl-tRNA synthetase. Nature Chemical Biology 8:311–317.

31. Matthews H, Usman-Idris M, Khan F, Read M, Nirmalan N. 2013. Drug repositioning as a route to anti-malarial drug discovery: preliminary investigation of the in vitro anti-malarial efficacy of emetine dihydrochloride hydrate. Malaria Journal 12:359.

32. Reynolds JM, El Bissati K, Brandenburg J, Günzl A, Mamoun CB. 2007. Antimalarial activity of the anticancer and proteasome inhibitor bortezomib and its analog ZL3B. BMC clinical pharmacology 7:13–13.

33. Crary JL, Haldar K. 1992. Brefeldin A inhibits protein secretion and parasite maturation in the ring stage of Plasmodium falciparum. Molecular and Biochemical Parasitology 53:185–192.

34. Andrews KT, Walduck A, Kelso MJ, Fairlie DP, Saul A, Parsons PG. 2000. Anti-malarial effect of histone deacetylation inhibitors and mammalian tumour cytodifferentiating agents. Int J Parasitol 30:761–8.

35. Fang P, Yu X, Jeong SJ, Mirando A, Chen K, Chen X, Kim S, Francklyn CS, Guo M. 2015. Structural basis for full-spectrum inhibition of translational functions on a tRNA synthetase. Nature Communications 6:6402.

36. Novoa EM, Camacho N, Tor A, Wilkinson B, Moss S, Marín-García P, Azcárate IG, Bautista JM, Mirando AC, Francklyn CS, Varon S, Royo M, Cortés A, Ribas de Pouplana L. 2014. Analogs of natural aminoacyl-tRNA synthetase inhibitors clear malaria in vivo. Proceedings of the National Academy of Sciences 111:E5508.

37. Herman JD, Pepper LR, Cortese JF, Estiu G, Galinsky K, Zuzarte-Luis V, Derbyshire ER, Ribacke U, Lukens AK, Santos SA, Patel V, Clish CB, Sullivan WJ, Jr., Zhou H, Bopp SE, Schimmel P, Lindquist S, Clardy J, Mota MM, Keller TL, Whitman M, Wiest O, Wirth DF, Mazitschek R. 2015. The cytoplasmic prolyl-tRNA synthetase of the malaria parasite is a dual-stage target of febrifugine and its analogs. Sci Transl Med 7:288ra77.

38. Wong W, Bai X-c, Brown A, Fernandez IS, Hanssen E, Condron M, Tan YH, Baum J, Scheres SHW. 2014. Cryo-EM structure of the Plasmodium falciparum 80S ribosome bound to the anti-protozoan drug emetine. eLife 3:e03080.

39. Liaoo L-L, Kupchan SM, Horwitz SB. 1976. Mode of Action of the Antitumor Compound Bruceantin, an Inhibitor of Protein Synthesis. Molecular Pharmacology 12:167.

40. Sridhar S, Bhat G, Guruprasad K. 2013. Analysis of bortezomib inhibitor docked within the catalytic subunits of the Plasmodium falciparum 20S proteasome. SpringerPlus 2:566–566.

41. Yoshida M, Kijima M, Akita M, Beppu T. 1990. Potent and specific inhibition of mammalian histone deacetylase both in vivo and in vitro by trichostatin A. Journal of Biological Chemistry 265:17174–17179.

42. Benting J, Mattei D, Lingelbach K. 1994. Brefeldin A inhibits transport of the glycophorin-binding protein from Plasmodium falciparum into the host erythrocyte. Biochem J 300 (Pt 3):821–6.

43. Hanson KK, Ressurreicao AS, Buchholz K, Prudencio M, Herman-Ornelas JD, Rebelo M, Beatty WL, Wirth DF, Hanscheid T, Moreira R, Marti M, Mota MM. 2013. Torins are potent antimalarials that block replenishment of Plasmodium liver stage parasitophorous vacuole membrane proteins. Proc Natl Acad Sci U S A 110:E2838–47.

44. Delves M, Plouffe D, Scheurer C, Meister S, Wittlin S, Winzeler EA, Sinden RE, Leroy D. 2012. The Activities of Current Antimalarial Drugs on the Life Cycle Stages of Plasmodium: A Comparative Study with Human and Rodent Parasites. PLOS Medicine 9:e1001169.

45. LaMonte GM, Rocamora F, Marapana DS, Gnädig NF, Ottilie S, Luth MR, Worgall TS, Goldgof GM, Mohunlal R, Santha Kumar TR, Thompson JK, Vigil E, Yang J, Hutson D, Johnson T, Huang J, Williams RM, Zou BY, Cheung AL, Kumar P, Egan TJ, Lee MCS, Siegel D, Cowman AF, Fidock DA, Winzeler EA. 2020. Pan-active imidazolopiperazine antimalarials target the Plasmodium falciparum intracellular secretory pathway. Nature Communications 11:1780.

46. LaMonte G, Lim MY-X, Wree M, Reimer C, Nachon M, Corey V, Gedeck P, Plouffe D, Du A, Figueroa N, Yeung B, Bifani P, Winzeler EA, Wellems TE. 2016. Mutations in the Plasmodium falciparum Cyclic Amine Resistance Locus (PfCARL) Confer Multidrug Resistance. mBio 7:e00696–16.

47. Nixon GL, Moss DM, Shone AE, Lalloo DG, Fisher N, O’Neill PM, Ward SA, Biagini GA. 2013. Antimalarial pharmacology and therapeutics of atovaquone. The Journal of antimicrobial chemotherapy 68:977–985.

48. Phillips MA, Lotharius J, Marsh K, White J, Dayan A, White KL, Njoroge JW, El Mazouni F, Lao Y, Kokkonda S, Tomchick DR, Deng X, Laird T, Bhatia SN, March S, Ng CL, Fidock DA, Wittlin S, Lafuente-Monasterio M, Benito FJG, Alonso LMS, Martinez MS, Jimenez-Diaz MB, Bazaga SF, Angulo-Barturen I, Haselden JN, Louttit J, Cui Y, Sridhar A, Zeeman A-M, Kocken C, Sauerwein R, Dechering K, Avery VM, Duffy S, Delves M, Sinden R, Ruecker A, Wickham KS, Rochford R, Gahagen J, Iyer L, Riccio E, Mirsalis J, Bathhurst I, Rueckle T, Ding X, Campo B, Leroy D, Rogers MJ, et al. 2015. A long-duration dihydroorotate dehydrogenase inhibitor (DSM265) for prevention and treatment of malaria. Science Translational Medicine 7:296ra111.

49. Paquet T, Manach CL, Cabrera DG, Younis Y, Henrich PP, Abraham TS, Lee MCS, Basak R, Ghidelli-Disse S, Lafuente-Monasterio MJ, Bantscheff M, Ruecker A, Blagborough AM, Zakutansky SE, Zeeman A-M, White KL, Shackleford DM, Mannila J, Morizzi J, Scheurer C, Angulo-Barturen I, Martínez MS, Ferrer S, Sanz LM, Gamo FJ, Reader J, Botha M, Dechering KJ, Sauerwein RW, Tungtaeng A, Vanachayangkul P, Lim CS, Burrows J, Witty MJ, Marsh KC, Bodenreider C, Rochford R, Solapure SM, Jiménez-Díaz MB, Wittlin S, Charman SA, Donini C, Campo B, Birkholtz L-M, Hanson KK, Drewes G, Kocken CHM, Delves MJ, Leroy D, Fidock DA, et al. 2017. Antimalarial efficacy of MMV390048, an inhibitor of *Plasmodium* phosphatidylinositol 4-kinase. Science Translational Medicine 9:eaad9735.

50. Falco EA, Goodwin LG, Hitchings GH, Rollo IM, Russell PB. 1951. 2:4-diaminopyrimidines-a new series of antimalarials. British journal of pharmacology and chemotherapy 6:185–200.

51. Yuthavong Y, Tarnchompoo B, Vilaivan T, Chitnumsub P, Kamchonwongpaisan S, Charman SA, McLennan DN, White KL, Vivas L, Bongard E, Thongphanchang C, Taweechai S, Vanichtanankul J, Rattanajak R, Arwon U, Fantauzzi P, Yuvaniyama J, Charman WN, Matthews D. 2012. Malarial dihydrofolate reductase as a paradigm for drug development against a resistance-compromised target. Proceedings of the National Academy of Sciences of the United States of America 109:16823–16828.

52. Hounkpatin AB, Kreidenweiss A, Held J. 2019. Clinical utility of tafenoquine in the prevention of relapse of Plasmodium vivax malaria: a review on the mode of action and emerging trial data. Infection and drug resistance 12:553–570.

53. Foley M, Tilley L. 1998. Quinoline Antimalarials: Mechanisms of Action and Resistance and Prospects for New Agents. Pharmacology & Therapeutics 79:55–87.

54. Wong W, Bai XC, Sleebs BE, Triglia T, Brown A, Thompson JK, Jackson KE, Hanssen E, Marapana DS, Fernandez IS, Ralph SA, Cowman AF, Scheres SHW, Baum J. 2017. Mefloquine targets the Plasmodium falciparum 80S ribosome to inhibit protein synthesis. Nat Microbiol 2:17031.

55. Rankin KE, Graewe S, Heussler VT, Stanway RR. 2010. Imaging liver-stage malaria parasites. Cellular Microbiology 12:569–579.

56. Leiriao P, Mota MM, Rodriguez A. 2005. Apoptotic Plasmodium-Infected Hepatocytes Provide Antigens to Liver Dendritic Cells. The Journal of Infectious Diseases 191:1576–1581.

57. Grollman AP, Walsh WttaoM. 1967. Inhibitors of Protein Biosynthesis: II. MODE OF ACTION OF ANISOMYCIN. Journal of Biological Chemistry 242:3226–3233.

58. Jackson KE, Habib S, Frugier M, Hoen R, Khan S, Pham JS, Ribas de Pouplana L, Royo M, Santos MA, Sharma A, Ralph SA. 2011. Protein translation in Plasmodium parasites. Trends Parasitol 27:467–76.

59. Hollien J. 2013. Evolution of the unfolded protein response. Biochimica et Biophysica Acta (BBA) - Molecular Cell Research 1833:2458–2463.

60. Meister S, Plouffe DM, Kuhen KL, Bonamy GMC, Wu T, Barnes SW, Bopp SE, Borboa R, Bright AT, Che J, Cohen S, Dharia NV, Gagaring K, Gettayacamin M, Gordon P, Groessl T, Kato N, Lee MCS, McNamara CW, Fidock DA, Nagle A, Nam T-g, Richmond W, Roland J, Rottmann M, Zhou B, Froissard P, Glynne RJ, Mazier D, Sattabongkot J, Schultz PG, Tuntland T, Walker JR, Zhou Y, Chatterjee A, Diagana TT, Winzeler EA. 2011. Imaging of Plasmodium liver stages to drive next- generation antimalarial drug discovery. Science (New York, NY) 334:1372–1377.

61. Kumar N, Koski G, Harada M, Aikawa M, Zheng H. 1991. Induction and localization of Plasmodium falciparum stress proteins related to the heat shock protein 70 family. Molecular and Biochemical Parasitology 48:47–58.

62. Kaiser G, De Niz M, Zuber B, Burda P-C, Kornmann B, Heussler VT, Stanway RR. 2016. High resolution microscopy reveals an unusual architecture of the Plasmodium berghei endoplasmic reticulum. Molecular Microbiology 102:775–791.

63. Gosline SJC, Nascimento M, McCall L-I, Zilberstein D, Thomas DY, Matlashewski G, Hallett M. 2011. Intracellular Eukaryotic Parasites Have a Distinct Unfolded Protein Response. PLOS ONE 6:e19118.

64. Ward P, Equinet L, Packer J, Doerig C. 2004. Protein kinases of the human malaria parasite Plasmodium falciparum: the kinome of a divergent eukaryote. BMC genomics 5:79–79.

65. Zhang M, Gallego-Delgado J, Fernandez-Arias C, Waters NC, Rodriguez A, Tsuji M, Wek RC, Nussenzweig V, Sullivan WJ, Jr. 2017. Inhibiting the Plasmodium eIF2alpha Kinase PK4 Prevents Artemisinin-Induced Latency. Cell Host Microbe 22:766–776 e4.

66. Bridgford JL, Xie SC, Cobbold SA, Pasaje CFA, Herrmann S, Yang T, Gillett DL, Dick LR, Ralph SA, Dogovski C, Spillman NJ, Tilley L. 2018. Artemisinin kills malaria parasites by damaging proteins and inhibiting the proteasome. Nat Commun 9:3801.

67. Zhang M, Mishra S, Sakthivel R, Rojas M, Ranjan R, Sullivan WJ, Fontoura BMA, Ménard R, Dever TE, Nussenzweig V. 2012. PK4, a eukaryotic initiation factor 2α(eIF2α) kinase, is essential for the development of the erythrocytic cycle of <em>Plasmodium</em>. Proceedings of the National Academy of Sciences 109:3956–3961.

68. Axten JM, Medina JR, Feng Y, Shu A, Romeril SP, Grant SW, Li WHH, Heerding DA, Minthorn E, Mencken T, Atkins C, Liu Q, Rabindran S, Kumar R, Hong X, Goetz A, Stanley T, Taylor JD, Sigethy SD, Tomberlin GH, Hassell AM, Kahler KM, Shewchuk LM, Gampe RT. 2012. Discovery of 7- Methyl-5-(1-{[3-(trifluoromethyl)phenyl]acetyl}-2,3-dihydro-1H-indol-5-yl)-7H-pyrrolo[2,3- d]pyrimidin-4-amine (GSK2606414), a Potent and Selective First-in-Class Inhibitor of Protein Kinase R (PKR)-like Endoplasmic Reticulum Kinase (PERK). Journal of Medicinal Chemistry 55:7193–7207.

69. Obeng EA, Carlson LM, Gutman DM, Harrington WJ, Jr., Lee KP, Boise LH. 2006. Proteasome inhibitors induce a terminal unfolded protein response in multiple myeloma cells. Blood 107:4907–16.

70. Van Voorhis WC, Adams JH, Adelfio R, Ahyong V, Akabas MH, Alano P, Alday A, Alemán Resto Y, Alsibaee A, Alzualde A, Andrews KT, Avery SV, Avery VM, Ayong L, Baker M, Baker S, Ben Mamoun C, Bhatia S, Bickle Q, Bounaadja L, Bowling T, Bosch J, Boucher LE, Boyom FF, Brea J, Brennan M, Burton A, Caffrey CR, Camarda G, Carrasquilla M, Carter D, Belen Cassera M, Chih- Chien Cheng K, Chindaudomsate W, Chubb A, Colon BL, Colón-López DD, Corbett Y, Crowther GJ, Cowan N, D’Alessandro S, Le Dang N, Delves M, DeRisi JL, Du AY, Duffy S, Abd El-Salam El-Sayed S, Ferdig MT, Fernández Robledo JA, Fidock DA, et al. 2016. Open Source Drug Discovery with the Malaria Box Compound Collection for Neglected Diseases and Beyond. PLOS Pathogens 12:e1005763.

71. Luth MR, Gupta P, Ottilie S, Winzeler EA. 2018. Using in Vitro Evolution and Whole Genome Analysis To Discover Next Generation Targets for Antimalarial Drug Discovery. ACS Infect Dis 4:301–314.

72. Okombo J, Kanai M, Deni I, Fidock DA. 2021. Genomic and Genetic Approaches to Studying Antimalarial Drug Resistance and Plasmodium Biology. Trends Parasitol 37:476–492.

73. Paquet T, Le Manach C, Cabrera DG, Younis Y, Henrich PP, Abraham TS, Lee MCS, Basak R, Ghidelli-Disse S, Lafuente-Monasterio MJ, Bantscheff M, Ruecker A, Blagborough AM, Zakutansky SE, Zeeman AM, White KL, Shackleford DM, Mannila J, Morizzi J, Scheurer C, Angulo-Barturen I, Martinez MS, Ferrer S, Sanz LM, Gamo FJ, Reader J, Botha M, Dechering KJ, Sauerwein RW, Tungtaeng A, Vanachayangkul P, Lim CS, Burrows J, Witty MJ, Marsh KC, Bodenreider C, Rochford R, Solapure SM, Jimenez-Diaz MB, Wittlin S, Charman SA, Donini C, Campo B, Birkholtz LM, Hanson KK, Drewes G, Kocken CHM, Delves MJ, Leroy D, Fidock DA, et al. 2017. Antimalarial efficacy of MMV390048, an inhibitor of Plasmodium phosphatidylinositol 4-kinase. Sci Transl Med 9.

74. Dziekan JM, Yu H, Chen D, Dai L, Wirjanata G, Larsson A, Prabhu N, Sobota RM, Bozdech Z, Nordlund P. 2019. Identifying purine nucleoside phosphorylase as the target of quinine using cellular thermal shift assay. Sci Transl Med 11.

75. Forte B, Ottilie S, Plater A, Campo B, Dechering KJ, Gamo FJ, Goldberg DE, Istvan ES, Lee M, Lukens AK, McNamara CW, Niles JC, Okombo J, Pasaje CFA, Siegel MG, Wirth D, Wyllie S, Fidock DA, Baragana B, Winzeler EA, Gilbert IH. 2021. Prioritization of Molecular Targets for Antimalarial Drug Discovery. ACS Infect Dis 7:2764–2776.

76. Lehane AM, Ridgway MC, Baker E, Kirk K. 2014. Diverse chemotypes disrupt ion homeostasis in the Malaria parasite. Mol Microbiol 94:327–39.

77. Allman EL, Painter HJ, Samra J, Carrasquilla M, Llinas M. 2016. Metabolomic Profiling of the Malaria Box Reveals Antimalarial Target Pathways. Antimicrob Agents Chemother 60:6635–6649.

78. Tamaki F, Fisher F, Milne R, Terán FS-R, Wiedemar N, Wrobel K, Edwards D, Baumann H, Gilbert IH, Baragana B, Baum J, Wyllie S. 2022. High-Throughput Screening Platform To Identify Inhibitors of Protein Synthesis with Potential for the Treatment of Malaria. Antimicrobial Agents and Chemotherapy 66:e00237–22.

79. Christophers SR, Fulton JD. 1939. Experiments with Isolated Malaria Parasites (Plasmodium Knowlesi) Free from Red Cells. Annals of Tropical Medicine & Parasitology 33:161–170.

80. Xie S, Griffin MDW, Winzeler EA, Ribas de Pouplana L, Tilley L. 2023. Targeting Aminoacyl tRNA Synthetases for Antimalarial Drug Development. Annu Rev Microbiol doi:10.1146/annurev-micro-032421-121210.

81. Ishiyama A, Iwatsuki M, Namatame M, Nishihara-Tsukashima A, Sunazuka T, Takahashi Y, Ōmura S, Otoguro K. 2011. Borrelidin, a potent antimalarial: stage-specific inhibition profile of synchronized cultures of Plasmodium falciparum. The Journal of Antibiotics 64:381–384.

82. Sugawara A, Tanaka T, Hirose T, Ishiyama A, Iwatsuki M, Takahashi Y, Otoguro K, Ōmura S, Sunazuka T. 2013. Borrelidin analogues with antimalarial activity: Design, synthesis and biological evaluation against Plasmodium falciparum parasites. Bioorganic & Medicinal Chemistry Letters 23:2302–2305.

83. Xie SC, Metcalfe RD, Dunn E, Morton CJ, Huang S-C, Puhalovich T, Du Y, Wittlin S, Nie S, Luth MR, Ma L, Kim M-S, Pasaje CFA, Kumpornsin K, Giannangelo C, Houghton FJ, Churchyard A, Famodimu MT, Barry DC, Gillett DL, Dey S, Kosasih CC, Newman W, Niles JC, Lee MCS, Baum J, Ottilie S, Winzeler EA, Creek DJ, Williamson N, Parker MW, Brand S, Langston SP, Dick LR, Griffin MDW, Gould AE, Tilley L. 2022. Reaction hijacking of tyrosine tRNA synthetase as a new whole-of-life-cycle antimalarial strategy. Science 376:1074–1079.

84. Bhatt TK, Kapil C, Khan S, Jairajpuri MA, Sharma V, Santoni D, Silvestrini F, Pizzi E, Sharma A. 2009. A genomic glimpse of aminoacyl-tRNA synthetases in malaria parasite Plasmodium falciparum. BMC genomics 10:644–644.

85. Jackson KE, Pham JS, Kwek M, De Silva NS, Allen SM, Goodman CD, McFadden GI, Ribas de Pouplana L, Ralph SA. 2012. Dual targeting of aminoacyl-tRNA synthetases to the apicoplast and cytosol in Plasmodium falciparum. International Journal for Parasitology 42:177–186.

86. Yu M, Kumar TRS, Nkrumah LJ, Coppi A, Retzlaff S, Li CD, Kelly BJ, Moura PA, Lakshmanan V, Freundlich JS, Valderramos J-C, Vilcheze C, Siedner M, Tsai JHC, Falkard B, Sidhu AbS, Purcell LA, Gratraud P, Kremer L, Waters AP, Schiehser G, Jacobus DP, Janse CJ, Ager A, Jacobs WR, Sacchettini JC, Heussler V, Sinnis P, Fidock DA. 2008. The Fatty Acid Biosynthesis Enzyme FabI Plays a Key Role in the Development of Liver-Stage Malarial Parasites. Cell Host & Microbe 4:567–578.

87. Caicedo JC, Cooper S, Heigwer F, Warchal S, Qiu P, Molnar C, Vasilevich AS, Barry JD, Bansal HS, Kraus O, Wawer M, Paavolainen L, Herrmann MD, Rohban M, Hung J, Hennig H, Concannon J, Smith I, Clemons PA, Singh S, Rees P, Horvath P, Linington RG, Carpenter AE. 2017. Data-analysis strategies for image-based cell profiling. Nat Methods 14:849–863.

88. Vembar SS, Droll D, Scherf A. 2016. Translational regulation in blood stages of the malaria parasite Plasmodium spp.: systems-wide studies pave the way. Wiley Interdiscip Rev RNA 7:772–792.

89. Choi J-Y, Kumar V, Pachikara N, Garg A, Lawres L, Toh JY, Voelker DR, Ben Mamoun C. 2016. Characterization of Plasmodium phosphatidylserine decarboxylase expressed in yeast and application for inhibitor screening. Molecular Microbiology 99:999–1014.

90. Subramanian G, Belekar MA, Shukla A, Tong JX, Sinha A, Chu TTT, Kulkarni AS, Preiser PR, Reddy DS, Tan KSW, Shanmugam D, Chandramohanadas R, Sullivan WJ. 2018. Targeted Phenotypic Screening in Plasmodium falciparum and Toxoplasma gondii Reveals Novel Modes of Action of Medicines for Malaria Venture Malaria Box Molecules. mSphere 3:e00534–17.

91. Plouffe David M, Wree M, Du Alan Y, Meister S, Li F, Patra K, Lubar A, Okitsu Shinji L, Flannery Erika L, Kato N, Tanaseichuk O, Comer E, Zhou B, Kuhen K, Zhou Y, Leroy D, Schreiber Stuart L, Scherer Christina A, Vinetz J, Winzeler Elizabeth A. 2016. High-Throughput Assay and Discovery of Small Molecules that Interrupt Malaria Transmission. Cell Host & Microbe 19:114–126.

92. Lucantoni L, Loganathan S, Avery VM. 2017. The need to compare: assessing the level of agreement of three high-throughput assays against Plasmodium falciparum mature gametocytes. Scientific Reports 7:45992.

93. Maher SP, Vantaux A, Chaumeau V, Chua ACY, Cooper CA, Andolina C, Péneau J, Rouillier M, Rizopoulos Z, Phal S, Piv E, Vong C, Phen S, Chhin C, Tat B, Ouk S, Doeurk B, Kim S, Suriyakan S, Kittiphanakun P, Awuku NA, Conway AJ, Jiang RHY, Russell B, Bifani P, Campo B, Nosten F, Witkowski B, Kyle DE. 2021. Probing the distinct chemosensitivity of Plasmodium vivax liver stage parasites and demonstration of 8-aminoquinoline radical cure activity in vitro. Scientific Reports 11:19905.

94. Allman EL, Painter HJ, Samra J, Carrasquilla M, Llinás M. 2016. Metabolomic Profiling of the Malaria Box Reveals Antimalarial Target Pathways. Antimicrobial Agents and Chemotherapy 60:6635–6649.

95. Istvan ES, Guerra F, Abraham M, Huang KS, Rocamora F, Zhao H, Xu L, Pasaje C, Kumpornsin K, Luth MR, Cui H, Yang T, Palomo Diaz S, Gomez-Lorenzo MG, Qahash T, Mittal N, Ottilie S, Niles J, Lee MCS, Llinas M, Kato N, Okombo J, Fidock DA, Schimmel P, Gamo FJ, Goldberg DE, Winzeler EA. 2023. Cytoplasmic isoleucyl tRNA synthetase as an attractive multistage antimalarial drug target. Sci Transl Med 15:eadc9249.

96. Prudencio M, Mota MM, Mendes AM. 2011. A toolbox to study liver stage malaria. Trends Parasitol 27:565–74.

97. Xie SC, Metcalfe RD, Dunn E, Morton CJ, Huang SC, Puhalovich T, Du Y, Wittlin S, Nie S, Luth MR, Ma L, Kim MS, Pasaje CFA, Kumpornsin K, Giannangelo C, Houghton FJ, Churchyard A, Famodimu MT, Barry DC, Gillett DL, Dey S, Kosasih CC, Newman W, Niles JC, Lee MCS, Baum J, Ottilie S, Winzeler EA, Creek DJ, Williamson N, Parker MW, Brand S, Langston SP, Dick LR, Griffin MDW, Gould AE, Tilley L. 2022. Reaction hijacking of tyrosine tRNA synthetase as a new whole-of-life-cycle antimalarial strategy. Science 376:1074–1079.

98. Valenciano AL, Gomez-Lorenzo MG, Vega-Rodriguez J, Adams JH, Roth A. 2022. In vitro models for human malaria: targeting the liver stage. Trends Parasitol 38:758–774.

99. Franke-Fayard B, Trueman H, Ramesar J, Mendoza J, van der Keur M, van der Linden R, Sinden RE, Waters AP, Janse CJ. 2004. A Plasmodium berghei reference line that constitutively expresses GFP at a high level throughout the complete life cycle. Mol Biochem Parasitol 137:23–33.

100. Tsuji M, Mattei D, Nussenzweig RS, Eichinger D, Zavala F. 1994. Demonstration of heat-shock protein 70 in the sporozoite stage of malaria parasites. Parasitology Research 80:16–21.

101. Trager W, Jensen JB. 1976. Human malaria parasites in continuous culture. Science 193:673–5.

102. Schindelin J, Arganda-Carreras I, Frise E, Kaynig V, Longair M, Pietzsch T, Preibisch S, Rueden C, Saalfeld S, Schmid B, Tinevez JY, White DJ, Hartenstein V, Eliceiri K, Tomancak P, Cardona A. 2012. Fiji: an open-source platform for biological-image analysis. Nat Methods 9:676–82.

103. Berthold MR, Cebron N, Dill F, Gabriel TR, Kötter T, Meinl T, Ohl P, Sieb C, Thiel K, Wiswedel B. KNIME: The Konstanz Information Miner, p 319–326. In (ed), Springer Berlin Heidelberg,

