## Supplementary Figures and Tables for "Single-cell quantitative bioimaging of *P. berghei* liver stage translation": McLellan et al Supplementary Figures.pdf

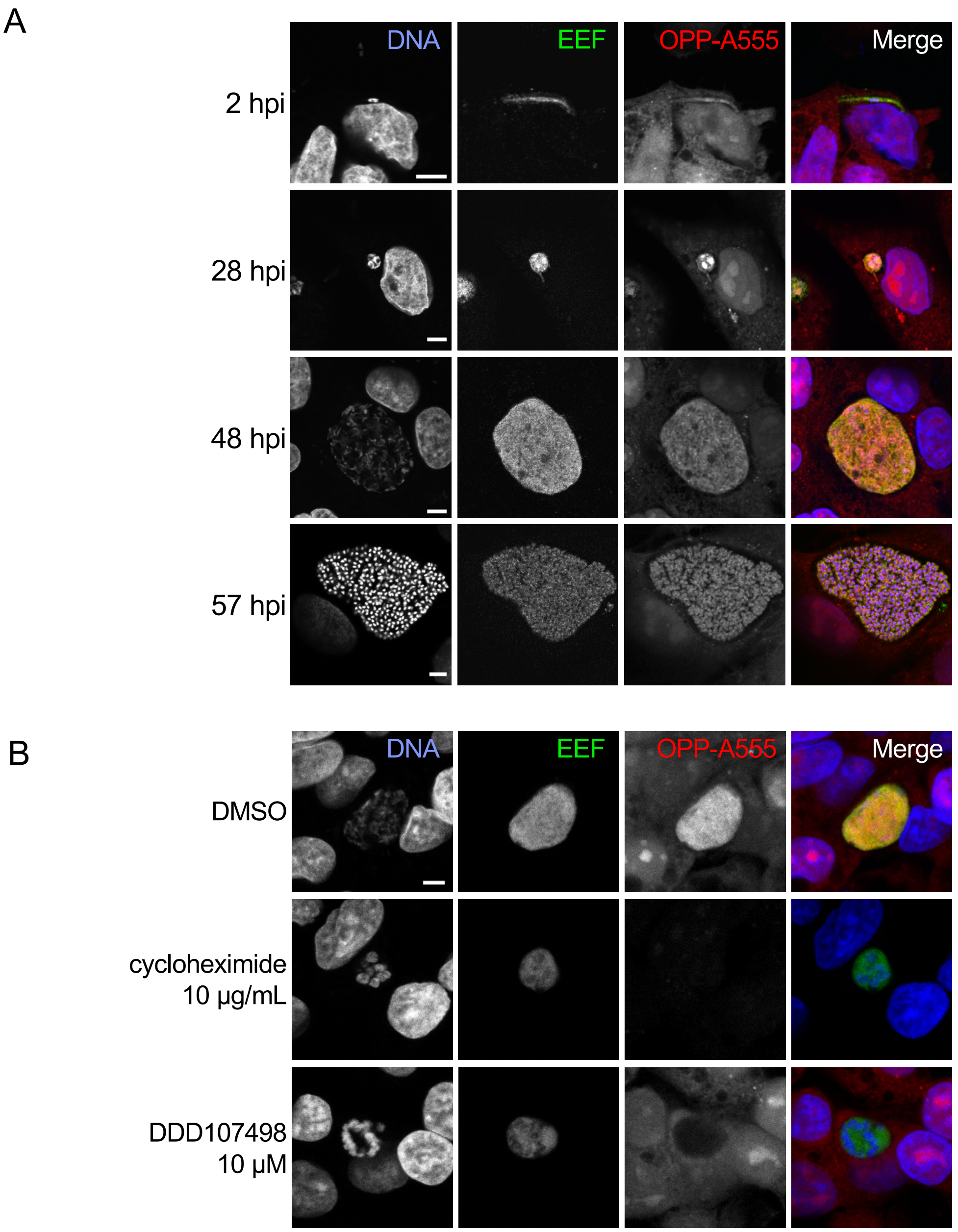


**Figure 1. Visualization of the nascent proteome in *Plasmodium berghei* liver stage parasites.** A-B) Representative, single confocal images of *P. berghei*-infected HepG2 cells, with OPP conjugated to Alexa Fluor 555 (OPP-A555) labeling the nascent proteome in both HepG2 and parasite (EEF), with Hoechst labeling DNA. Single channel images are shown in grayscale, with merges pseudo colored as labeled. A) Visualization of the nascent proteome throughout liver stage development, with parasite immunolabeled with α-UIS4 (2 hpi) or α-HSP70 (28, 48, and 57 hpi). B) Nascent proteome visualized in infected HepG2 cells following treatment from 44-48 hpi with translation inhibitors cycloheximide (10 µg/mL), or DDD107498 (10 µM) vs. DMSO control. All images in B) were acquired and processed with identical settings. Scale bars = 5 µm.

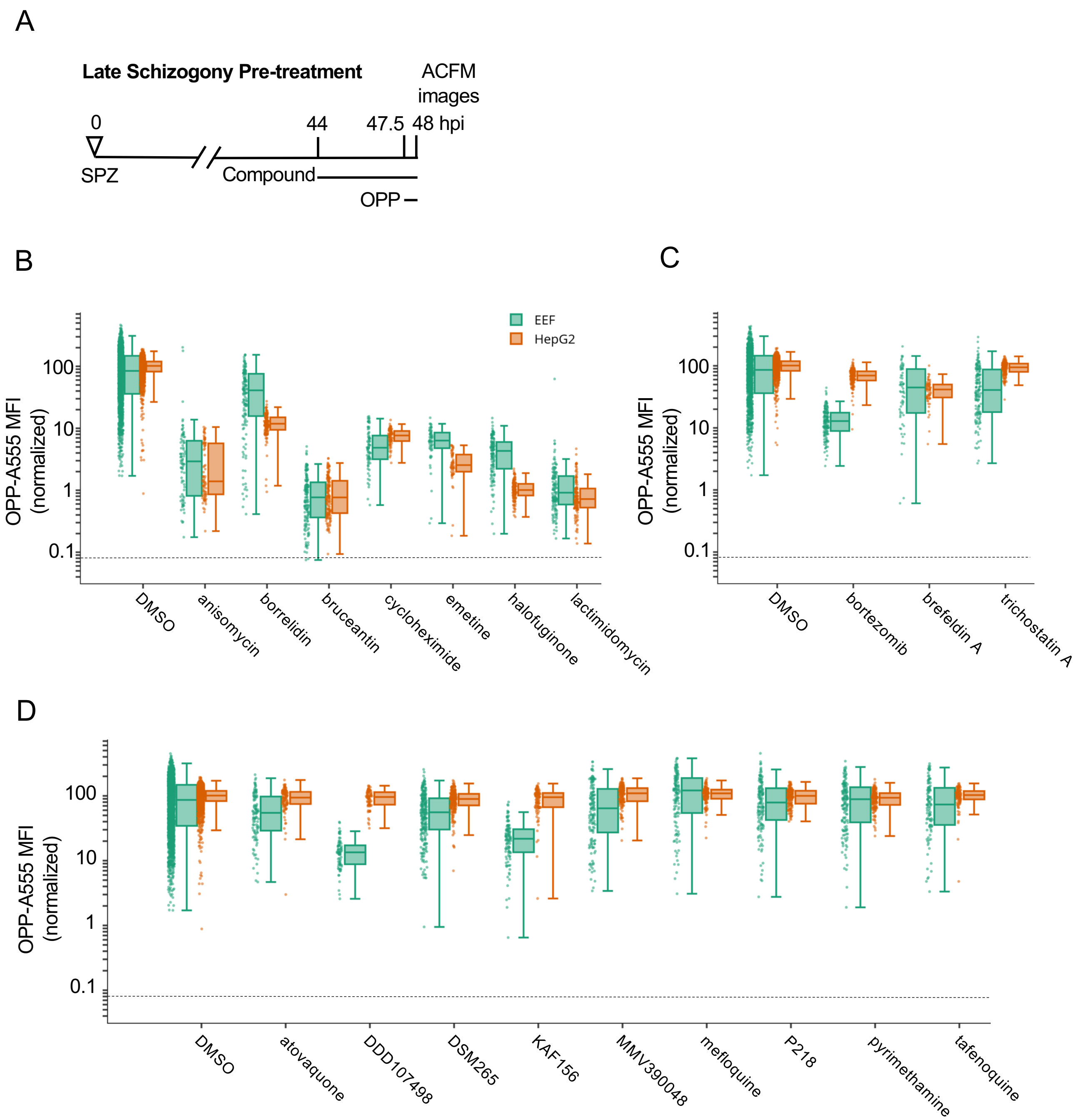


**Figure 2. Testing select bioactive compounds for inhibition of *Plasmodium* liver stage translation.** Quantification of *P. berghei* and HepG2 protein synthesis after 3.5h pre-treatment with diverse, active compounds, as schematized in A). See Table S-1 for compound details. B-D) Boxplots quantifying translation inhibition via OPP-A555 mean fluorescence intensity (OPP-A555 MFI) from all single parasite ACFM images acquired for n≥3 independent experiments, with each dot corresponding to a single EEF (green) or associated in-image HepG2 cells (orange). Specific signal cutoff (see Figure S2-3) is indicated by the dashed line. Compounds tested are known pan-eukaryotic translation inhibitors B), pan-eukaryotic inhibitors of cellular processes other than translation C), and antimalarial compounds D). Compounds were tested at 10 µM except for cycloheximide (10 µg/mL), tafenoquine (1.25 µM), mefloquine (2.5 µM), trichostatin A (5 µM) & brefeldin A (5 µg/mL).

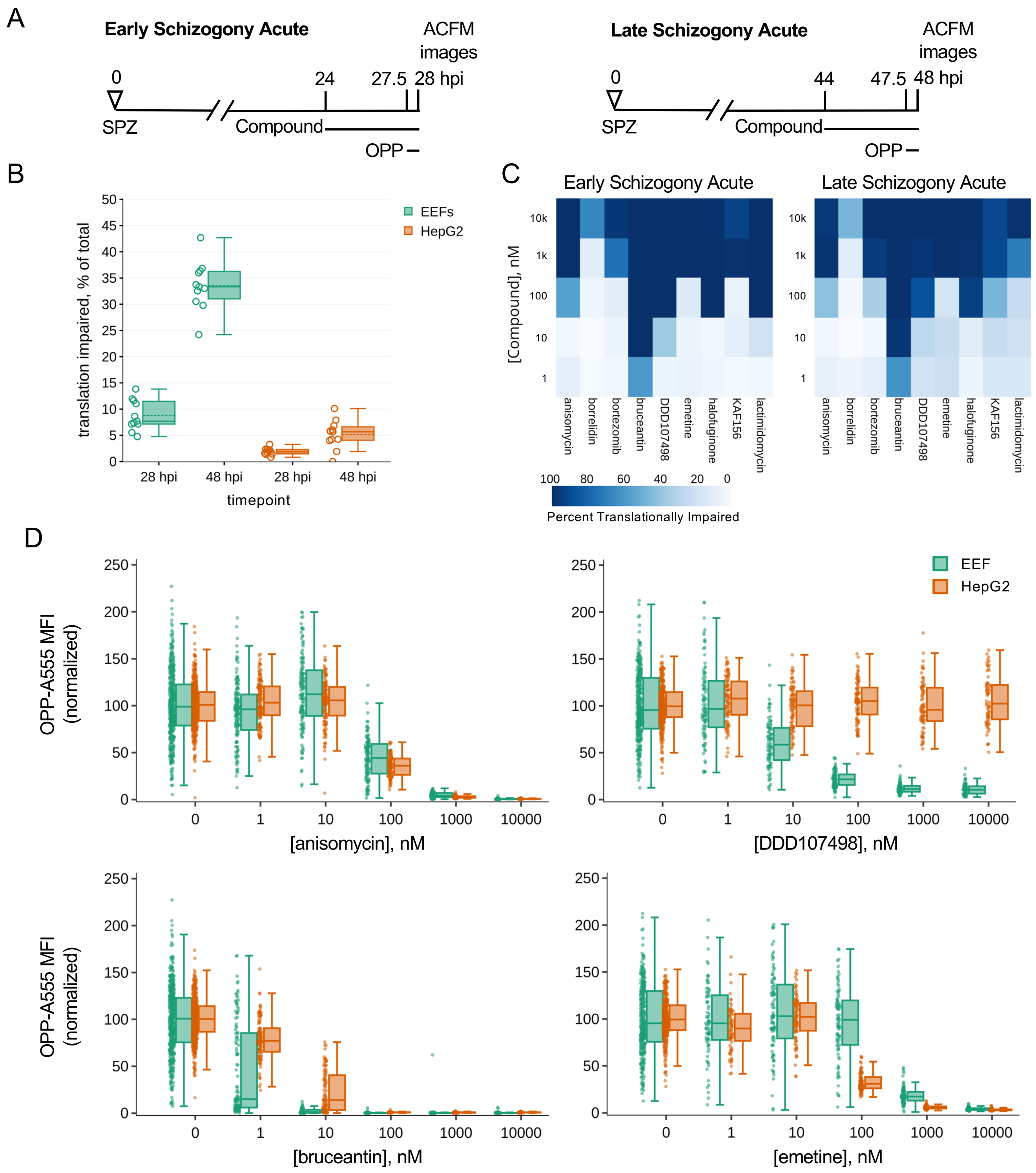


**Figure 3. Assessing heterogeneity and potency of translation inhibition during early and late schizogony.** A) Experimental schematics. B) DMSO treated control EEFs and corresponding in-image HepG2 cells were classed as translationally impaired (individual parasite OPP-A555 MFI ≤ 50% of experiment OPP-A555 mean) or unimpaired during early and late schizogony; data show the mean of 11 matched independent experiments with circles representing individual experiment values. C-D) Determining potency of translation inhibition in *P. berghei* and in-image HepG2 cells after acute pre-treatment with the inhibitors identified in Fig. 2; n≥3 independent experiments C) Percentage of single EEFs categorized as translationally impaired for inhibitors across concentrations in early and late schizogony. D) Concentration-response data in single parasites and in-image HepG2 cells for select translation inhibitors; all data normalized to mean of in-plate DMSO controls, set to 100.

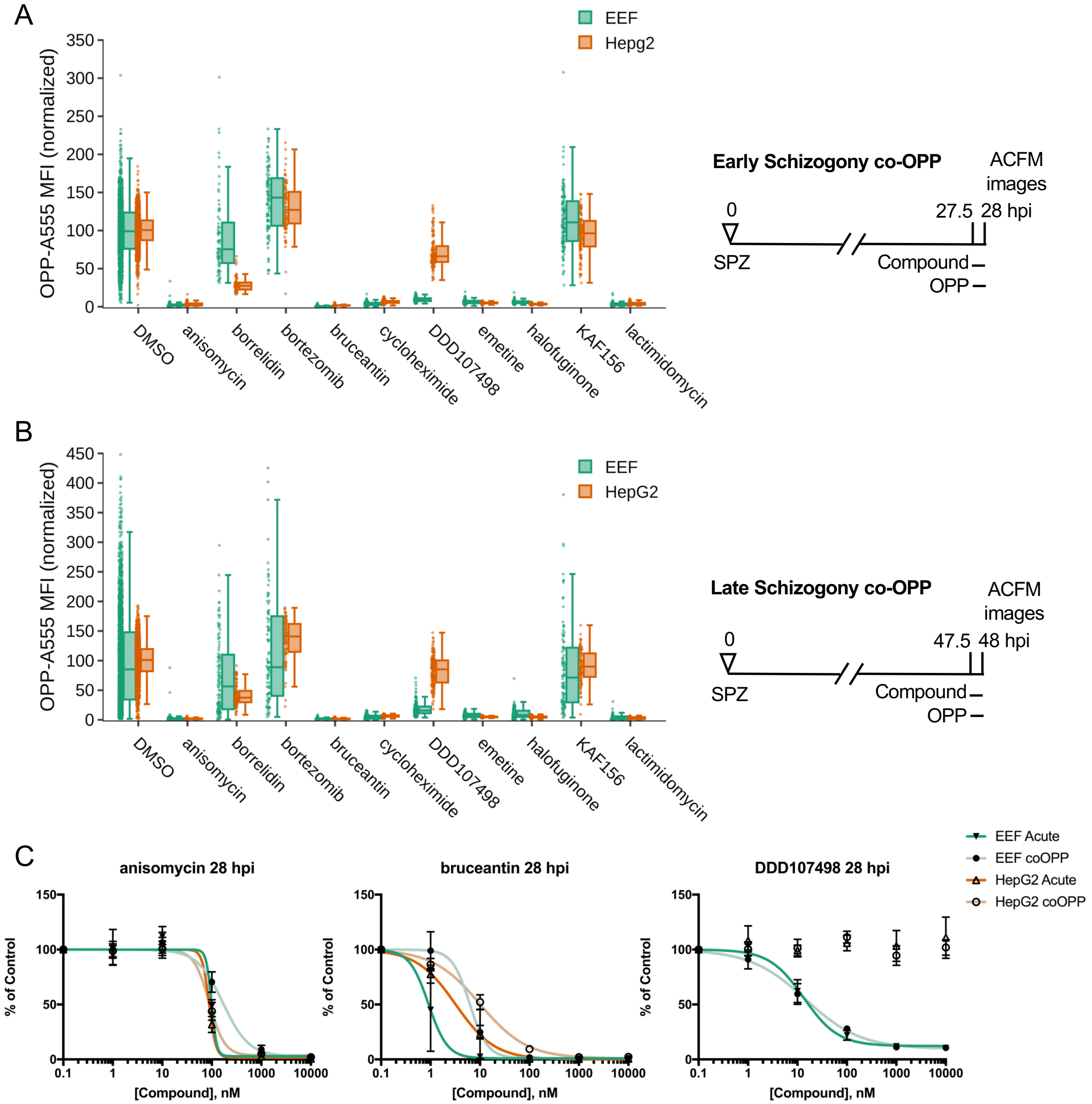


EEF $EC_{50}$ acute: 99 nM
EEF $EC_{50}$ coOPP: 168 nM
HepG2 $EC_{50}$ acute: 88 nM
HepG2 $EC_{50}$ coOPP: 89 nM

EEF $EC_{50}$ acute: 0.9 nM
EEF $EC_{50}$ coOPP: 6.2 nM
HepG2 $EC_{50}$ acute: 3.2 nM
HepG2 $EC_{50}$ coOPP: 10 nM

EEF $EC_{50}$ acute: 13 nM
EEF $EC_{50}$ coOPP: 15 nM
HepG2 $EC_{50}$ acute: na
HepG2 $EC_{50}$ coOPP: na

**Figure 4. Identification of direct vs. indirect translation inhibitors.** Quantification of protein synthesis in *P. berghei*-infected HepG2 cells in which active compounds at maximal concentrations were added together with OPP in early (A) and late (B) schizogony as described in figure schematics. Compound concentrations tested are the same as in Fig. 2. Each data point represents the normalized OPP-A555 mean fluorescence intensity (OPP-A555 MFI) of a single EEF or the corresponding HepG2 cells as labeled. C) Comparing coOPP and acute pre-treatment (from Fig. S3-1) concentration-response curves. All data shown was collected in n ≥ 3 independent experiments.

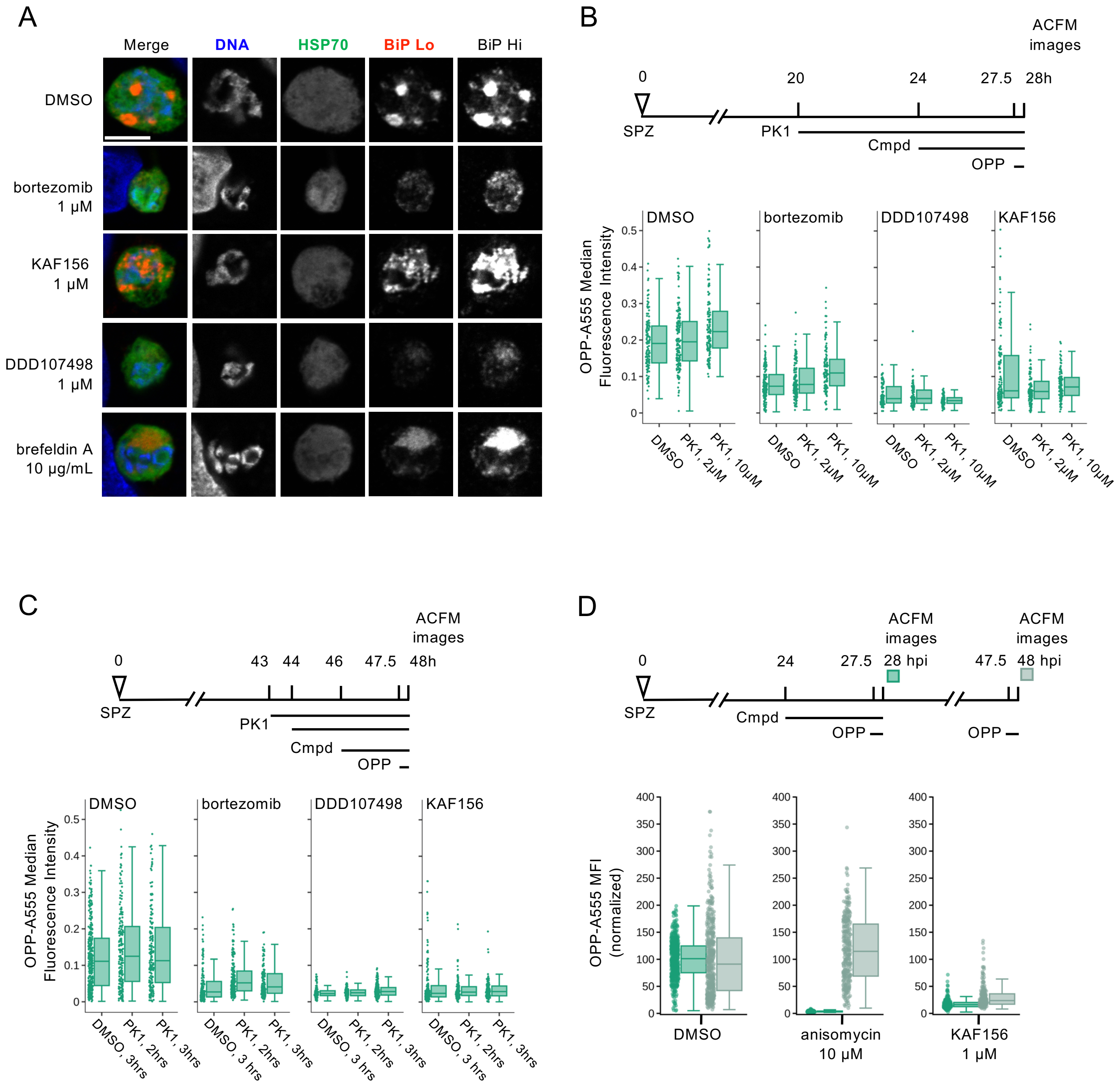


**Figure 5. Investigating the mechanism behind indirect translation inhibition**. A) Representative, single confocal images of *P. berghei* liver stage ER morphology after 4h compound treatment in early schizogony, at 28 hpi. Single channel images, all acquired with identical settings, are shown in grayscale, with merges pseudocolored as labeled; HSP70 marks the parasite and BiP specifically labels the parasite ER. Two images of BiP immunofluorescence were acquired with different gains (BiP Lo and BiP Hi) to visualize ER morphology across the range of BiP intensity observed. Scale bar = 5μm. B-D) Quantification of protein synthesis in single EEFs following treatments detailed in associated schematics. In B-C) [bortezomib] = 1 μM, [KAF156] = 0.5 μM, and [DDD107498] = 0.1 μM were used to achieve similar levels of submaximal translational inhibition in the parasites. n≥3 independent experiments. [PK1] as labelled in B), and 20 μM in C). Data in D) was normalized to the mean of the DMSO control parasites for each timepoint.

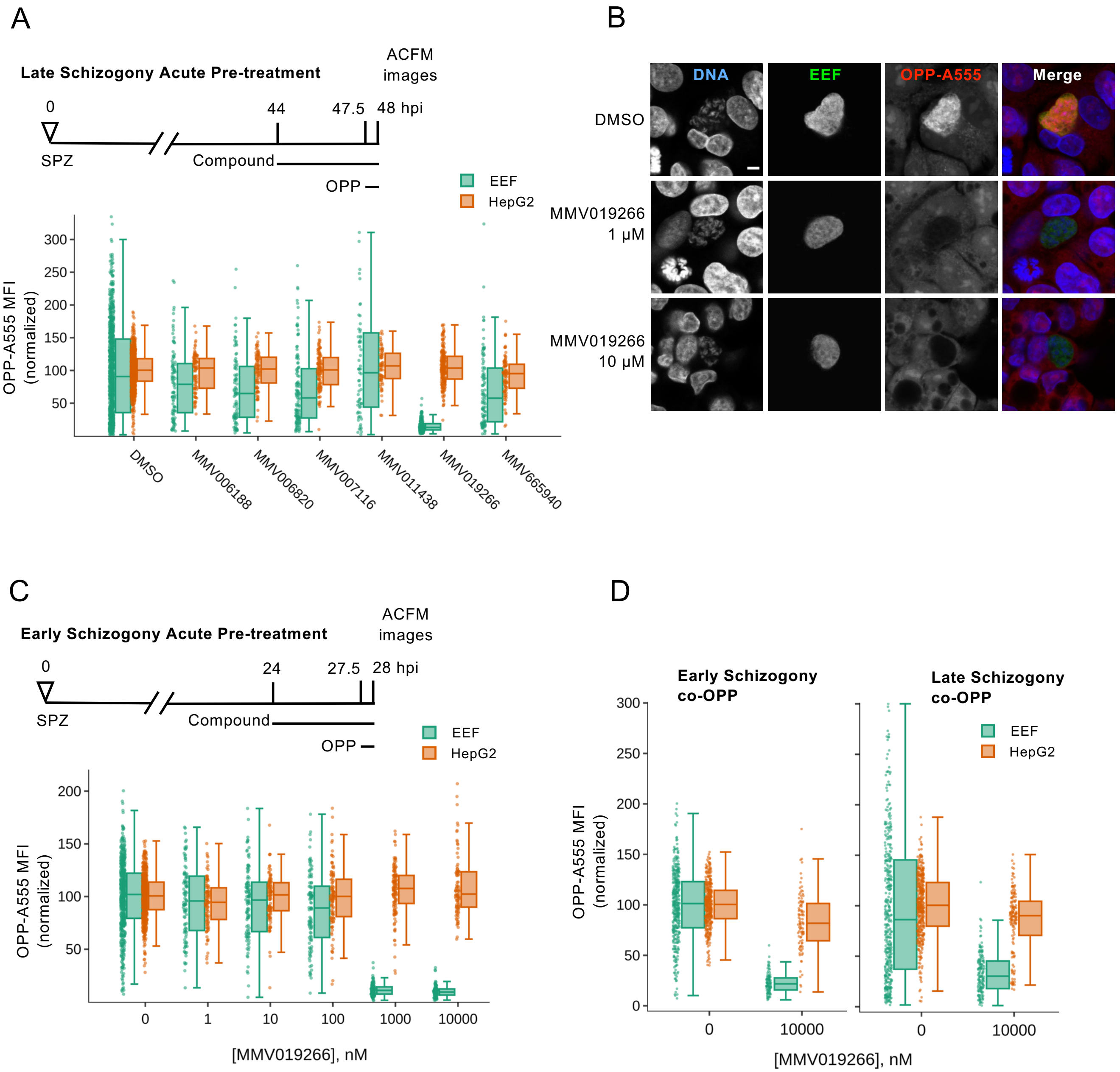


**Figure 6. Characterization of MMV019266 inhibition of *P. berghei* liver stage translation.** A) Select liver stage actives from the Malaria Box were tested at 10 μM for ability to inhibit *P. berghei* liver stage translation following acute pre-treatment in late schizogony. B) Representative single confocal images of OPP-A55 labeling after 4h acute pre-treatments with MMV019266 vs. control in late schizogony; merges are pseudo colored as indicated with parasite (EEF) immunolabeled with α-HSP70, and DNA stained with Hoechst. Scale bar = 5 μm. C-D) Quantification of protein synthesis in *P. berghei*-infected HepG2 cells; compound treatments as described. Each data point represents OPP-A555 mean fluorescence intensity (MFI) normalized to in-plate controls; n≥3 independent experiments.

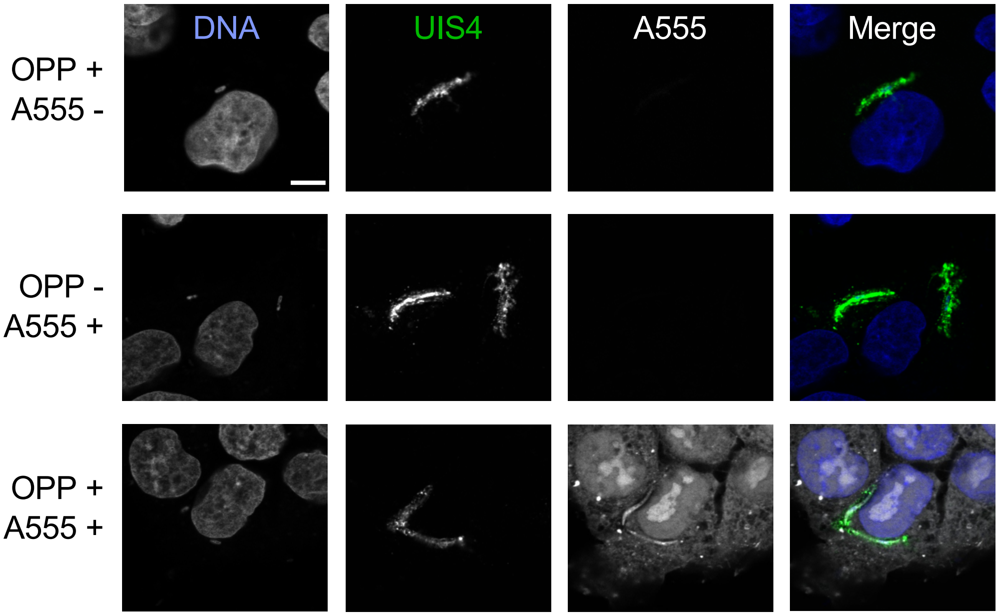


**Figure S1. Specificity of the OPP-A555 nascent proteome signal.** Control experiments were performed to visualize the specificity of OPP-A555 labeling of *P. berghei*-infected HepG2 at 2 hpi. Infected cells were incubated with (OPP+) or without (OPP-) OPP for 30 minutes before fixation. Following fixation, click labeling reactions were performed with (AF555+) and without (AF555-) addition of fluorophore to the labeling reaction mix. Representative, single confocal images were acquired and processed with identical settings. Single channel images are shown in grayscale, merged images are pseudo colored as labeled, with α-UIS4 marking the parasite and Hoechst staining DNA. Scale bar = 5 μm.

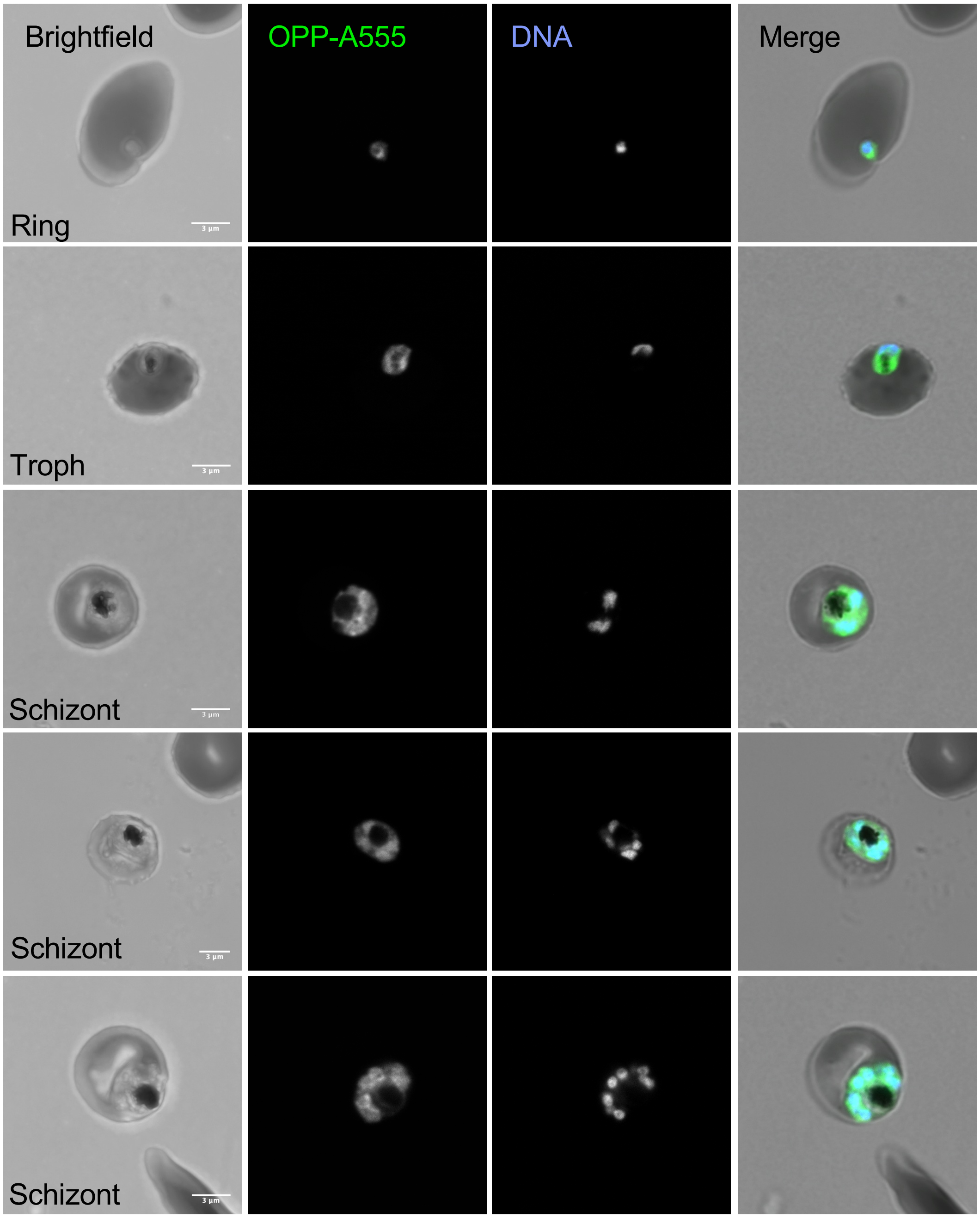


**Figure S2. Visualization of the nascent proteome in *Plasmodium falciparum* asexual blood stage parasites**. Representative single confocal images (fluorescence and merge) and maximum intensity projections (brightfield) of *Plasmodium falciparum*-infected erythrocytes with OPP-A555 labeling the nascent proteome and Hoechst labeling DNA. Scale bar = 3 μm.

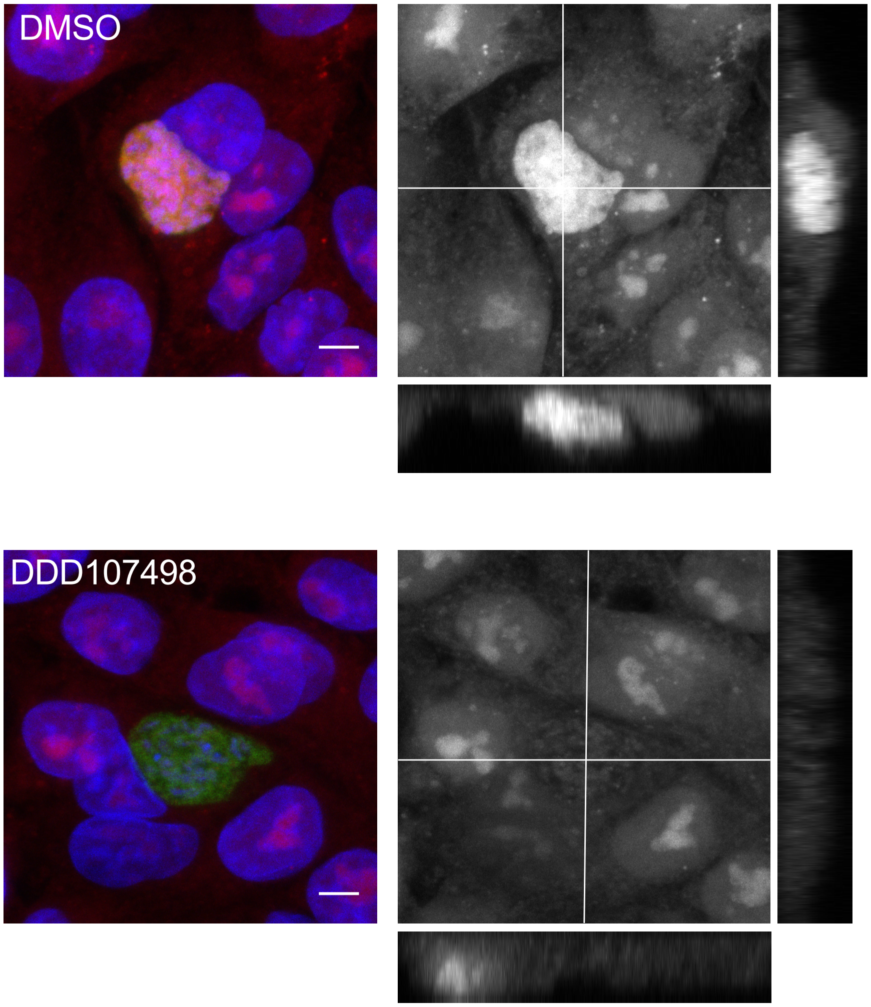


**Figure S3. OPP-A555 fluorescence in the host hepatocyte does not preclude quantification of the parasite signal in confocal images.** Confocal z-stacks through *P. berghei*-infected HepG2 cells with OPP-A555 labeling the nascent proteome following acute pre-treatment, are shown in maximum intensity projections in grayscale (xy) flanked by xz and yz confocal stacks at the positions indicated by intersecting lines in the xy projection. Image stacks are comprised of 11 (DMSO) and 9 (DDD107498) confocal images acquired with a 1μm z-step. Merged images are pseudo colored with OPP-A555 labeling the nascent proteome in red, Hoechst staining DNA in blue, and α-PbHSP70 immunolabeling the EEF in green. The xy images shown illustrate the the slice that would be algorithmically chosen for ACFM imaging to maximize the intensity of the PbHSP70 signal. Scale bars = 5 μm.

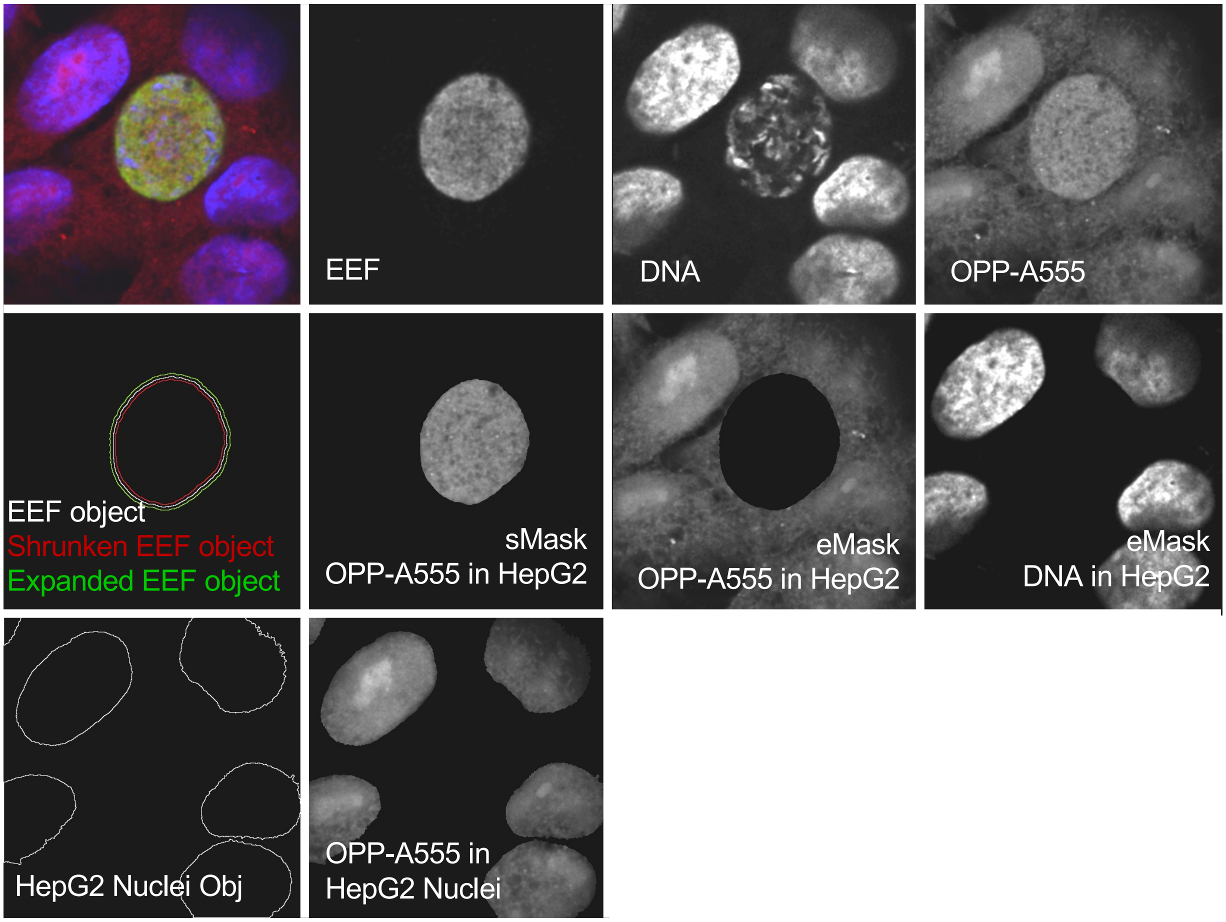


**Figure S4. Overview of image segmentation and masking for specific quantification of host and parasite protein synthesis.** After ACFM image acquisition, the EEF image (anti-HSP70) was segmented in CellProfiler to identify the set of labeled pixels, defined as the EEF object. The EEF object is then computationally shrunk and expanded by 2 pixels to generate the shrunken EEF and expanded EEF objects respectively, which exclude the host-parasite interface pixels. The shrunken EEF object is used to mask DNA and OPP-A555 (sMask as shown) for further segmentation and/or quantification of parasite features. The expanded EEF object is used as an inverted mask (eMask as shown) to select all pixels outside the parasite, allowing identification of HepG2 nuclei objects, and quantification of features inside the HepG2 cells.

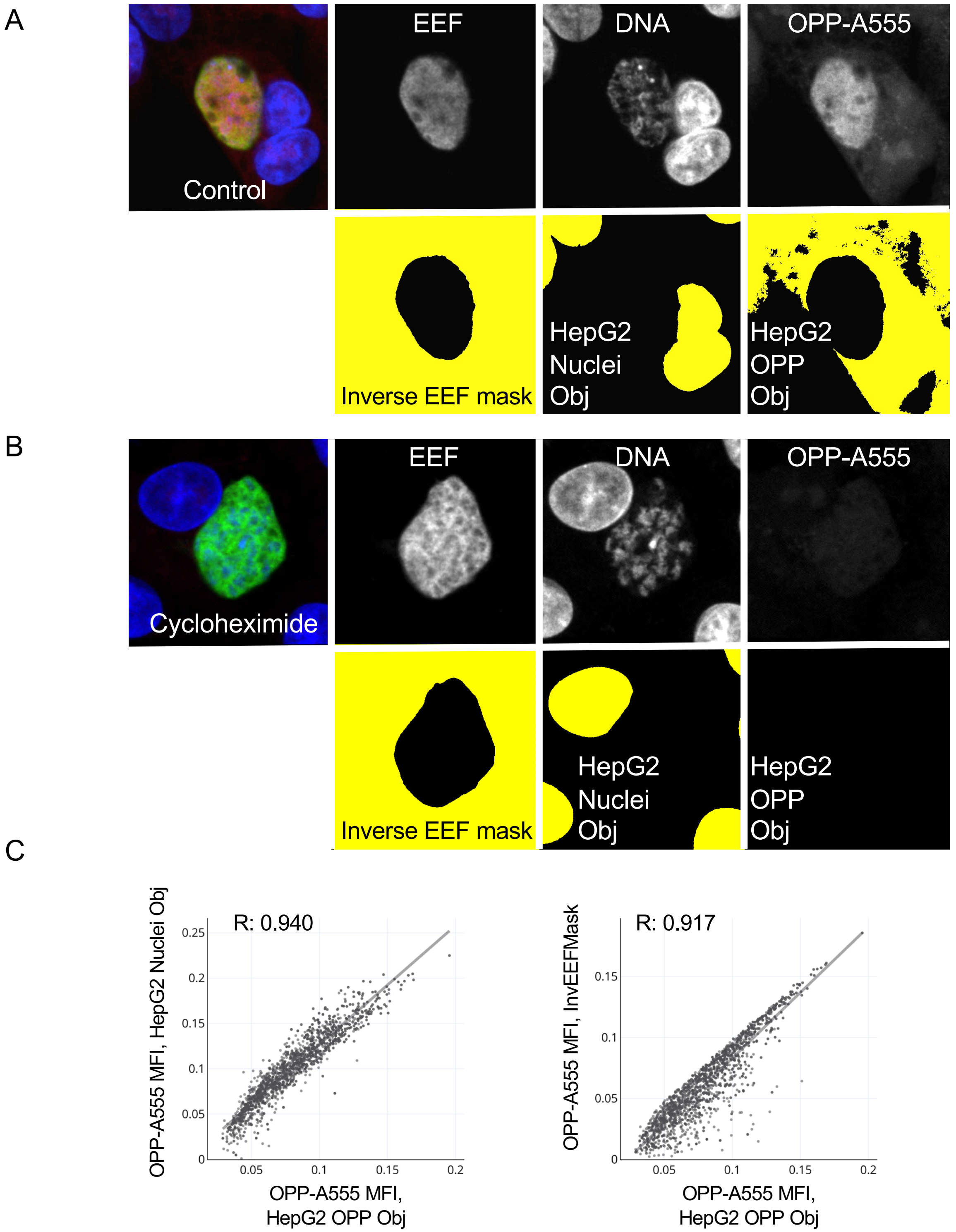


**Figure S5. The OPP-A555 signal in HepG2 nuclei is well correlated with whole cell translation intensity.** HepG2 protein synthesis was quantified in confocal ACFM images of *P. berghei*-infected HepG2 cells using three different strategies. Pixels marked for quantification are pseudo colored yellow in A-B). While the actual HepG2 OPP-A555 signal can be easily segmented (HepG2 OPP Obj) control cells (A), this method fails for images where HepG2 translation has been blocked by an inhibitor such as cycloheximide (B). Two additional strategies not dependent on segmentation of the OPP-A555 signal also implemented for comparison to the translation intensity of the HepG2 OPP Obj: quantification of OPP-A555 signal in all pixels of the inverse EEF mask (InvEEFMask) and in the segmented HepG2 nuclei (HepG2 Nuclei Obj), as shown in A-B). C) Correlation of OPP-A555 MFI from the HepG2 OPP Obj with that of HepG2 Nuclei Obj and InvEEFMask using a control parasite dataset consisting of 3779 single parasite ACFM images (single dot) from 13 independent infections, all with 44-48 hpi DMSO treatment, OPP labeling, and fixation at 48 hpi. R (correlation) is indicated for each.

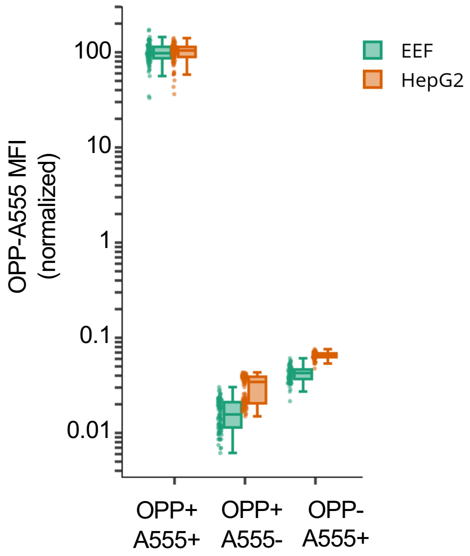


**Figure S6. Quantifying signal specificity for OPP-A55 labeling of the nascent proteome.** *P. berghei* -infected HepG2 cells were labeled with OPP (OPP+) or DMSO (OPP-) for 30 minutes before fixation at 28 hpi. Following fixation, click reactions were performed with (A555+) and without (A555-) addition of the fluorophore to the labeling reaction mix, followed by immunolabeling of parasites and DNA staining. Images were acquired with ACFM using the OPP+ A555+ to determine acquisition settings used for all conditions, and the mean of this condition was set to 100 and used to normalize all data. Boxplots quantify the amount of signal detected with and without the OPP and fluorophore addition, with each point corresponding to a single parasite, or the associated in-image HepG2 cells. Data is from a single experiment.

A

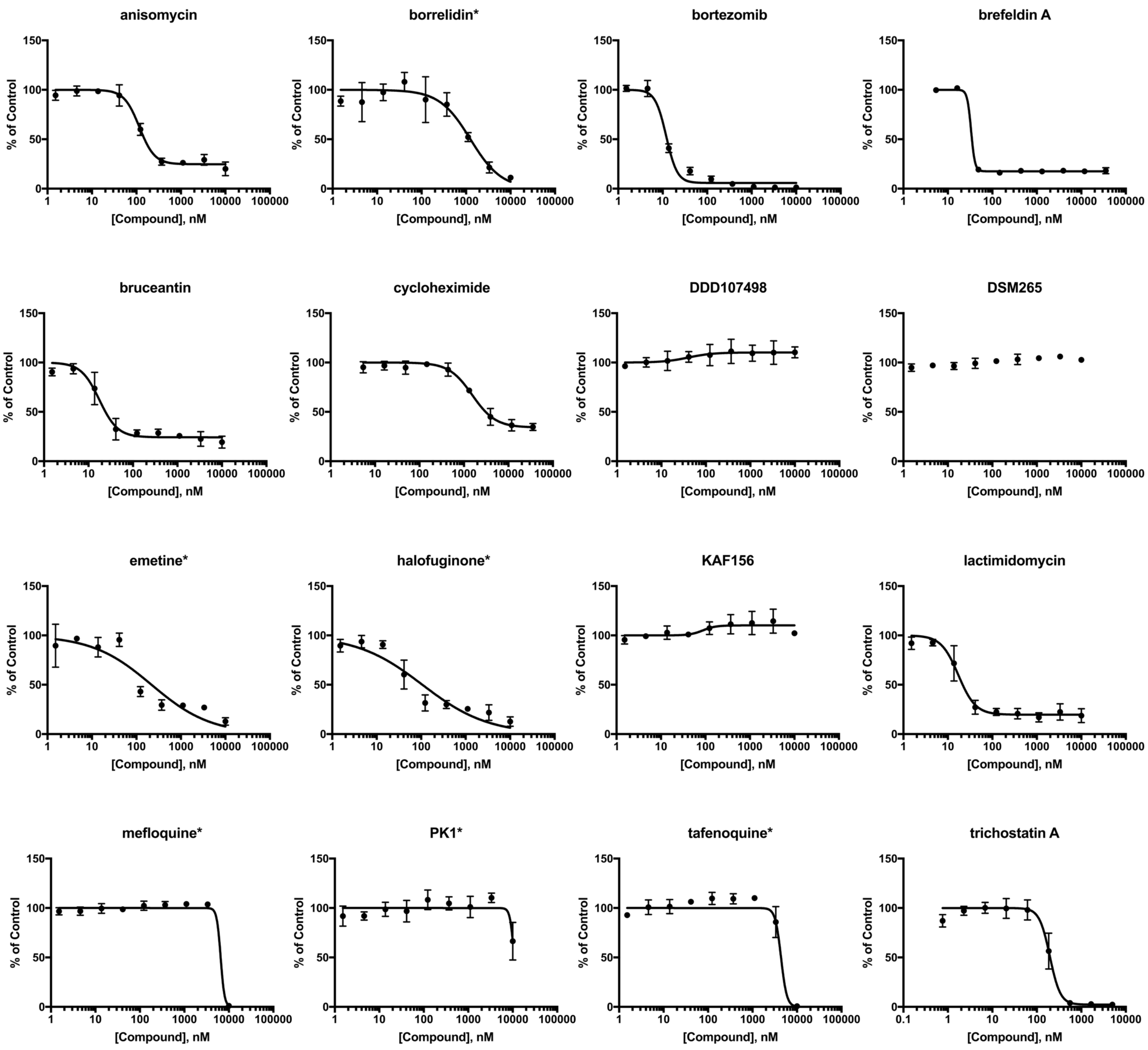


**Figure S7. HepG2 cytotoxicity of compounds tested for ability to inhibit liver stage translation.** Cellular viability of uninfected HepG2 cells determined by AlamarBlue fluorescence following a 48h compound treatment with 3-fold serial dilution; starting maximal concentration 10μM, except cycloheximide (10μg/mL). Concentration-response curves marked with an asterisk were fit with the bottom of the curve constrained to 0, while all others were fit open as detailed in Materials and Methods. Points represent the mean of 3 independent experiments, and errors bars show standard deviation.

Figure S13 Fig. S3-3

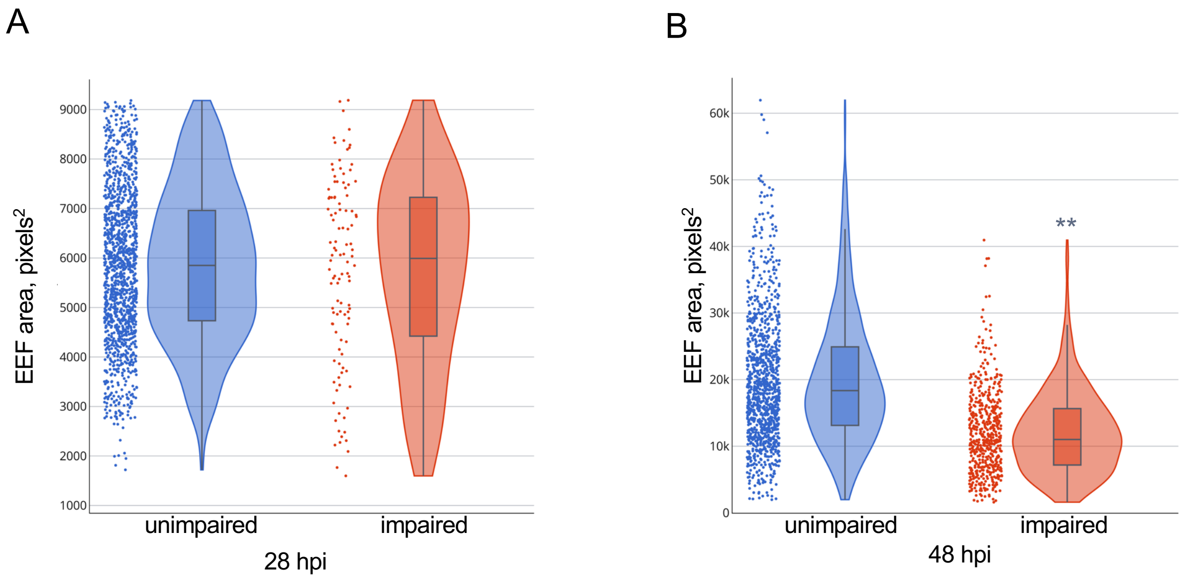


**Figure S8. Quantifying the relationship between parasite size and translation intensity in control EEFs.** A-B) Area of single, DMSO-treated control parasites, classed as translationally impaired or unimpaired (Fig. 3B dataset, see Fig. 3 legend for classification definition) at 28- and 48 hpi. To ensure that only images containing a single EEF were analyzed at 28 hpi, parasites with an area larger than the $9^{th}$ decile for the entire dataset in Fig. A (9188 pixels$^2$) were excluded from the analysis. No filtering was applied to the 48 hpi data, as wide variation in parasite size exists at this timepoint. Paired, two-tailed t-tests were run on mean area (unnormalized) of EEFs assigned to either translation class, from a total of 11 matched independent experiments. ** = $p < 0.005$.

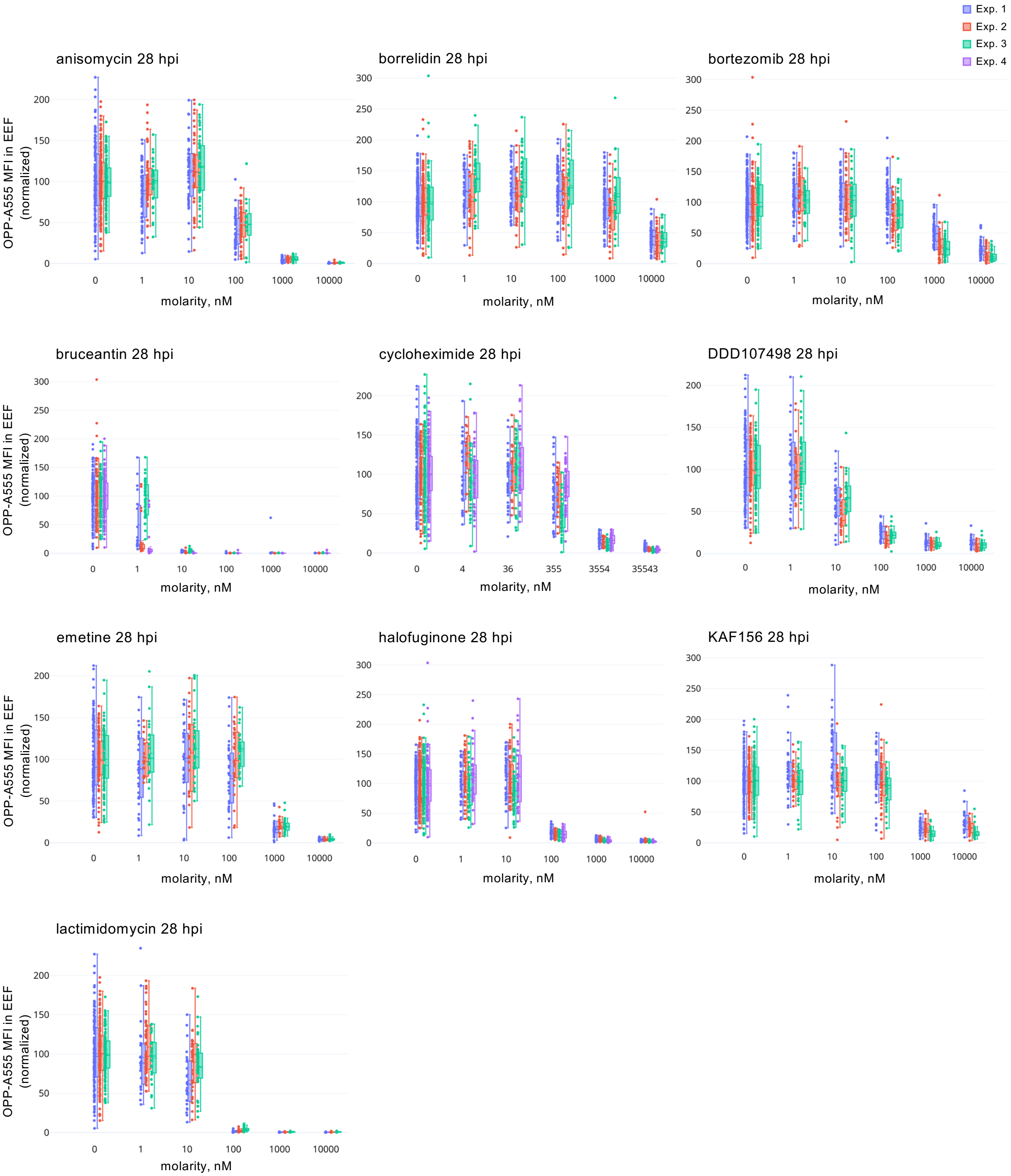
Exp. 1
Exp. 2
Exp. 3
Exp. 4
anisomycin 28 hpi
borrelidin 28 hpi
bortezomib 28 hpi
bruceantin 28 hpi
cycloheximide 28 hpi
DDD107498 28 hpi
emetine 28 hpi
halofuginone 28 hpi
KAF156 28 hpi
lactimidomycin 28 hpi
OPP-A555 MFI in EEF (normalized)
molarity, nM
0
1
10
100
1000
10000
4
36
355
3554
35543

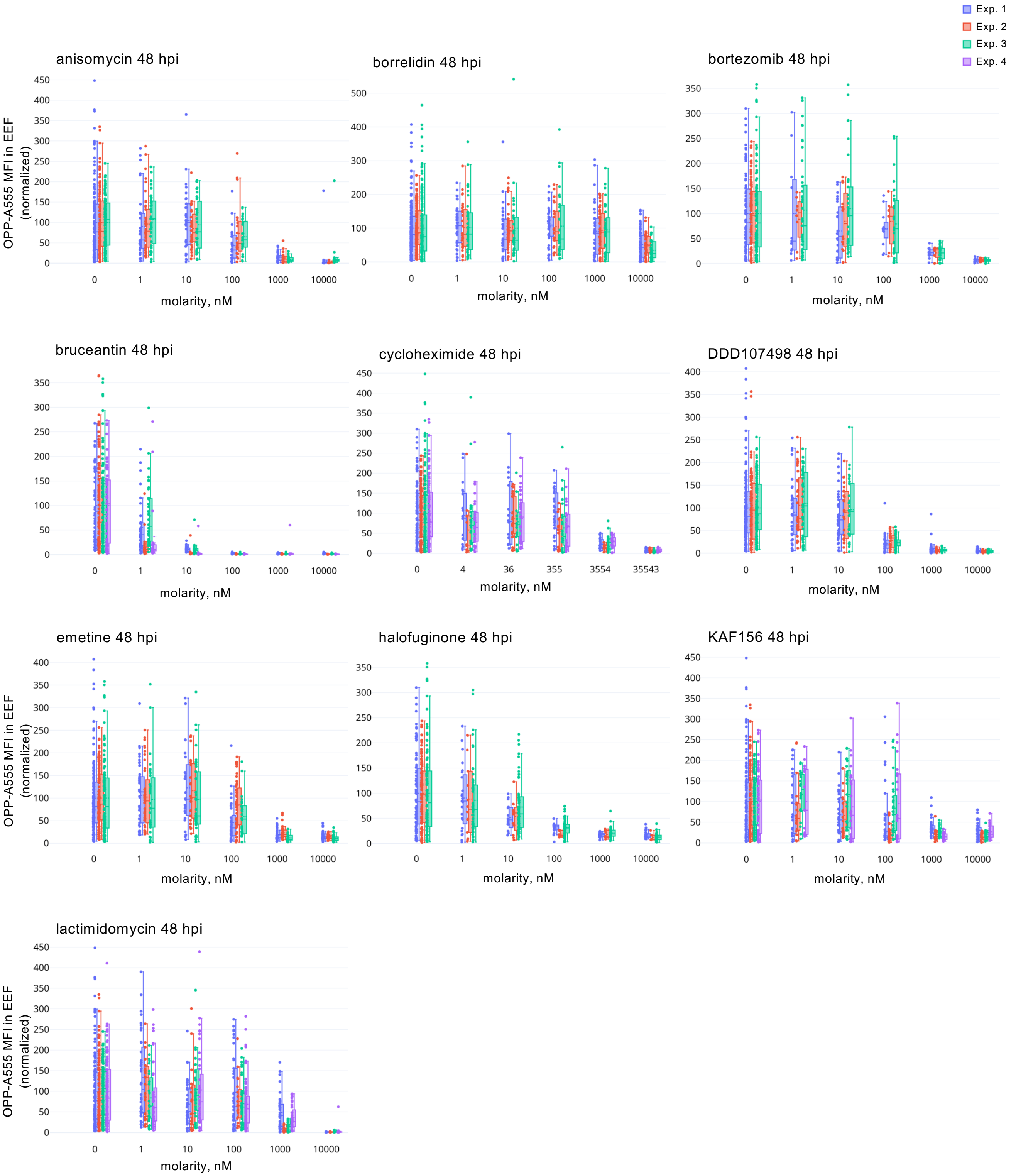


**Figure S9. Concentration dependent translation inhibition in EEFs by experiment.** Protein synthesis (OPP-A555 MFI) in single EEFs was quantified and normalized to in-plate DMSO controls following acute pre-treatment with a 5-point, 10-fold serial dilution with timepoint and compound treatment as labeled. Same data shown in Fig. 3C; here, data for each independent experiment is (n≥3) shown separately.

A

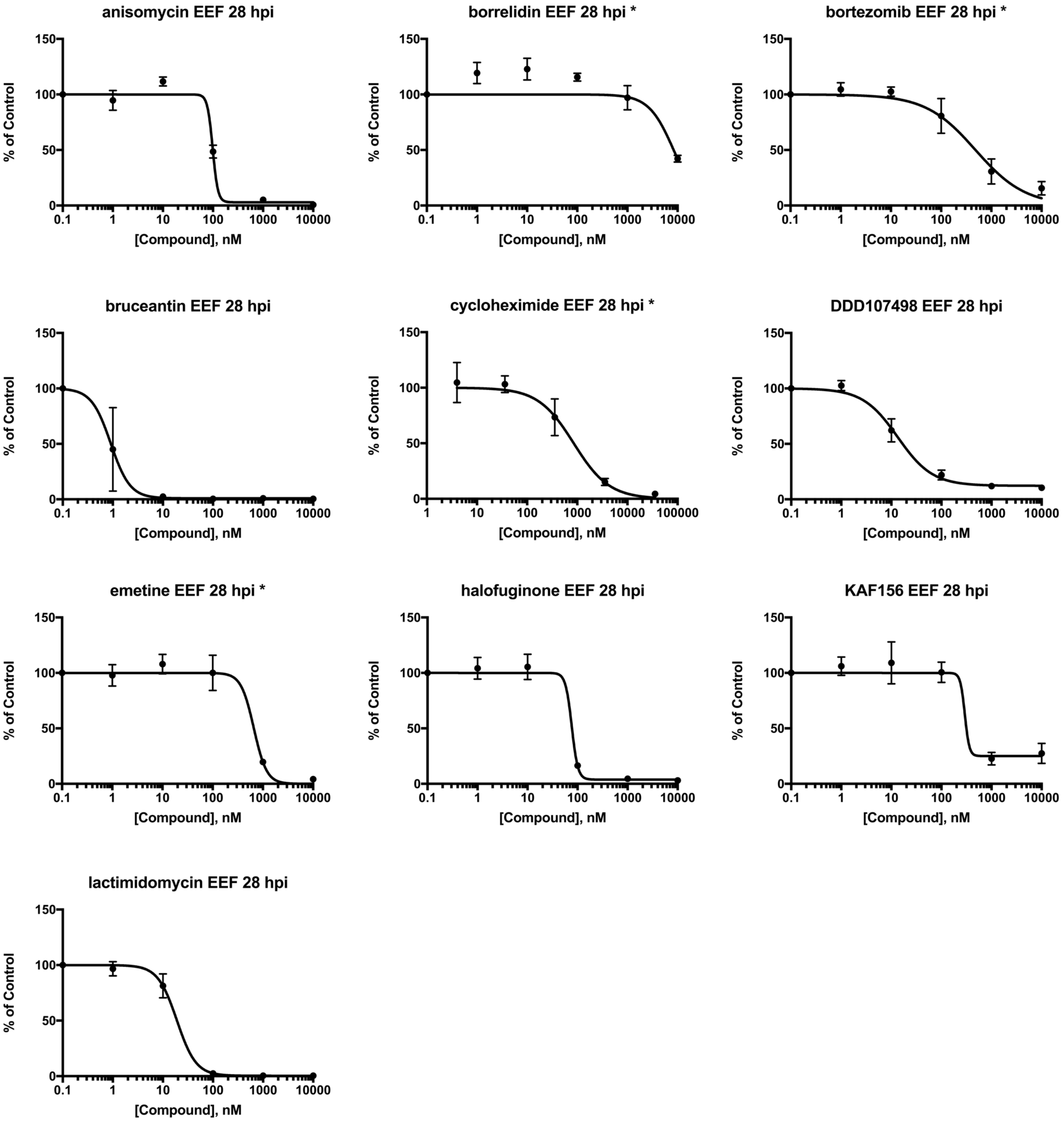

anisomycin EEF 28 hpi
borrelidin EEF 28 hpi *
bortezomib EEF 28 hpi *
bruceantin EEF 28 hpi
cycloheximide EEF 28 hpi *
DDD107498 EEF 28 hpi
emetine EEF 28 hpi *
halofuginone EEF 28 hpi
KAF156 EEF 28 hpi
lactimidomycin EEF 28 hpi
% of Control
0
50
100
150
[Compound], nM
0.1
1
10
100
1000
10000
100000

B

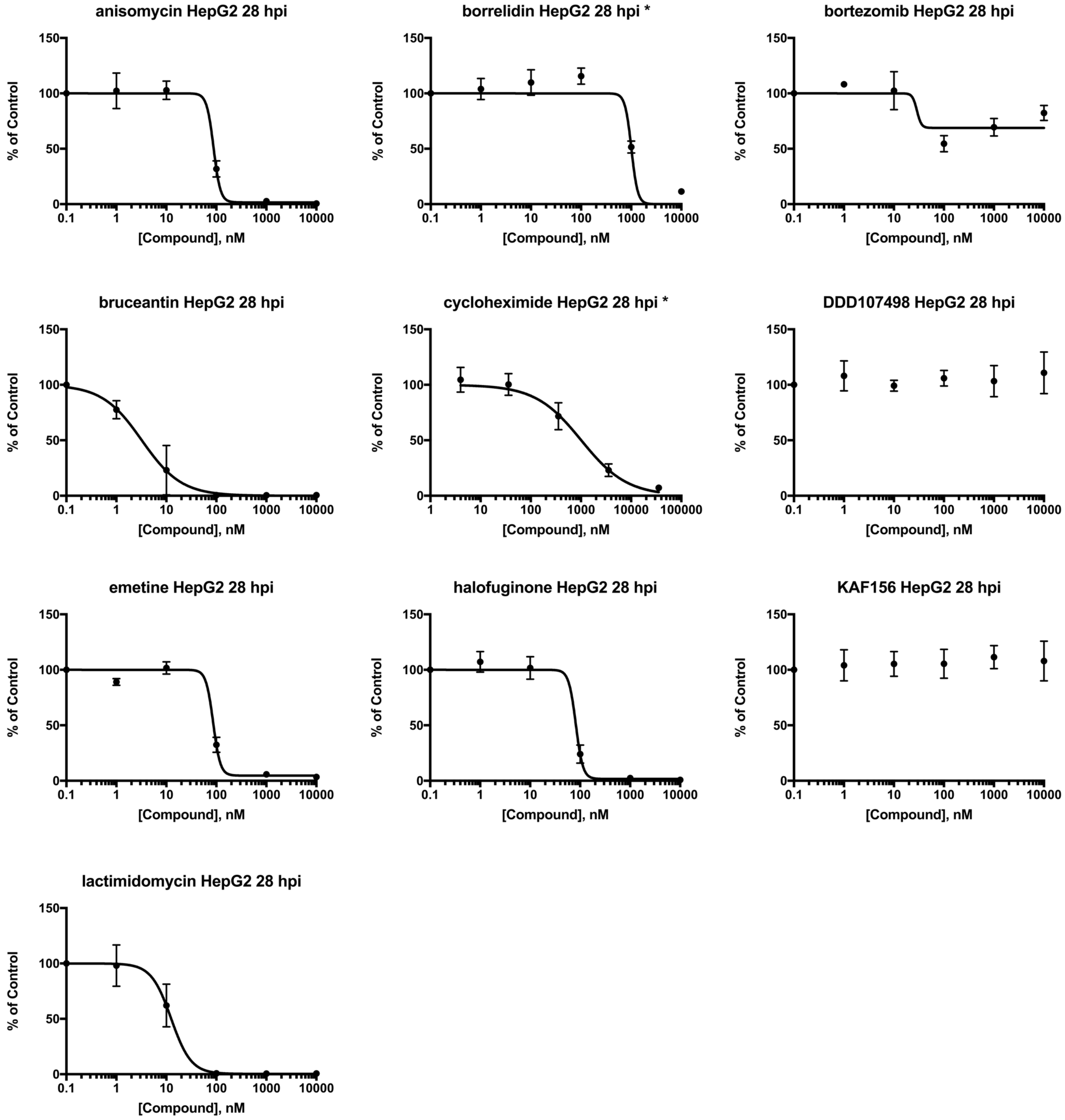
anisomycin HepG2 28 hpi
borrelidin HepG2 28 hpi *
bortezomib HepG2 28 hpi
bruceantin HepG2 28 hpi
cycloheximide HepG2 28 hpi *
DDD107498 HepG2 28 hpi
emetine HepG2 28 hpi
halofuginone HepG2 28 hpi
KAF156 HepG2 28 hpi
lactimidomycin HepG2 28 hpi
% of Control
[Compound], nM
0
50
100
150
0.1
1
10
100
1000
10000
100000

C

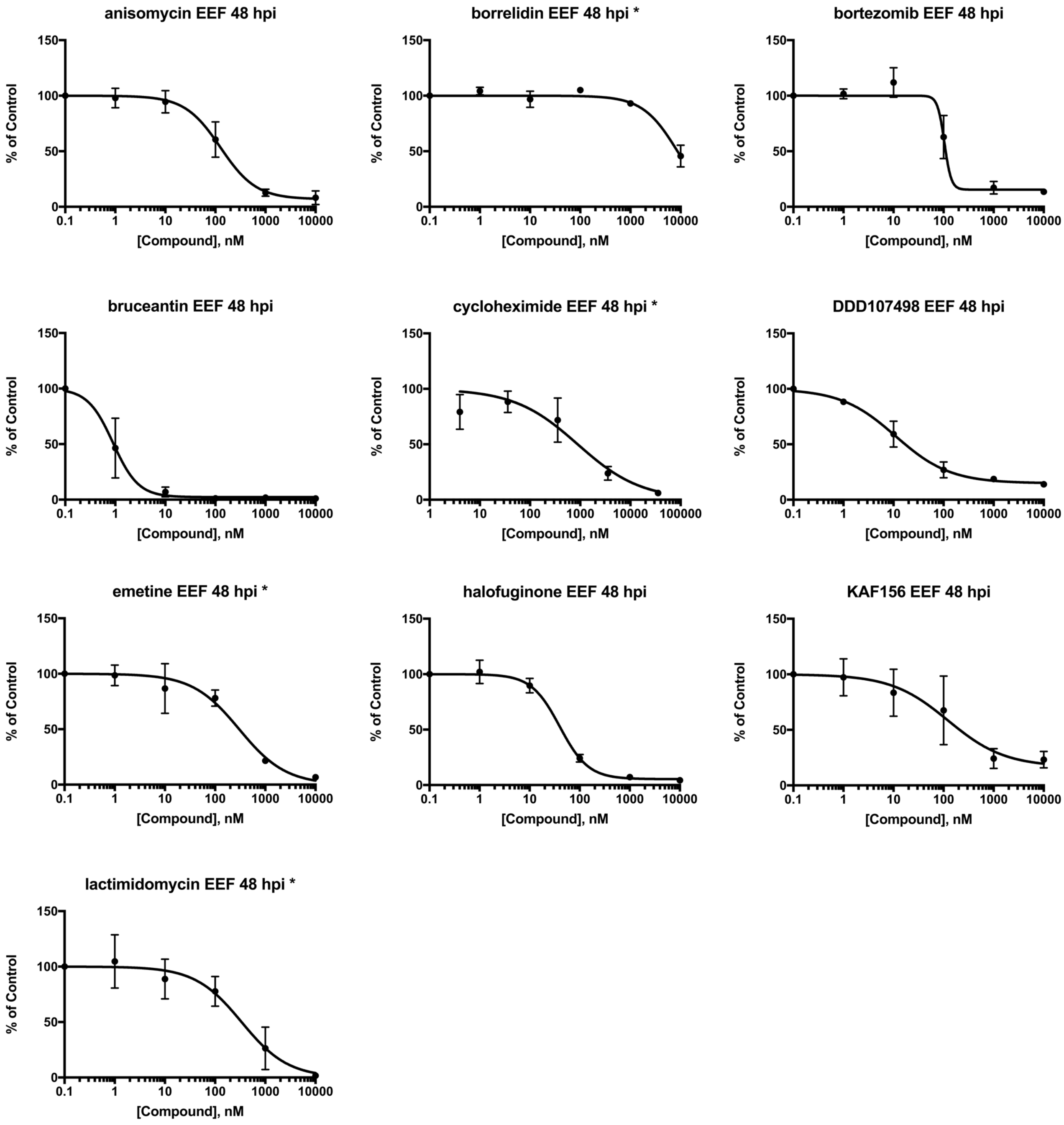
anisomycin EEF 48 hpi
borrelidin EEF 48 hpi *
bortezomib EEF 48 hpi
bruceantin EEF 48 hpi
cycloheximide EEF 48 hpi *
DDD107498 EEF 48 hpi
emetine EEF 48 hpi *
halofuginone EEF 48 hpi
KAF156 EEF 48 hpi
lactimidomycin EEF 48 hpi *
% of Control
[Compound], nM
0
50
100
150
0.1
1
10
100
1000
10000
100000

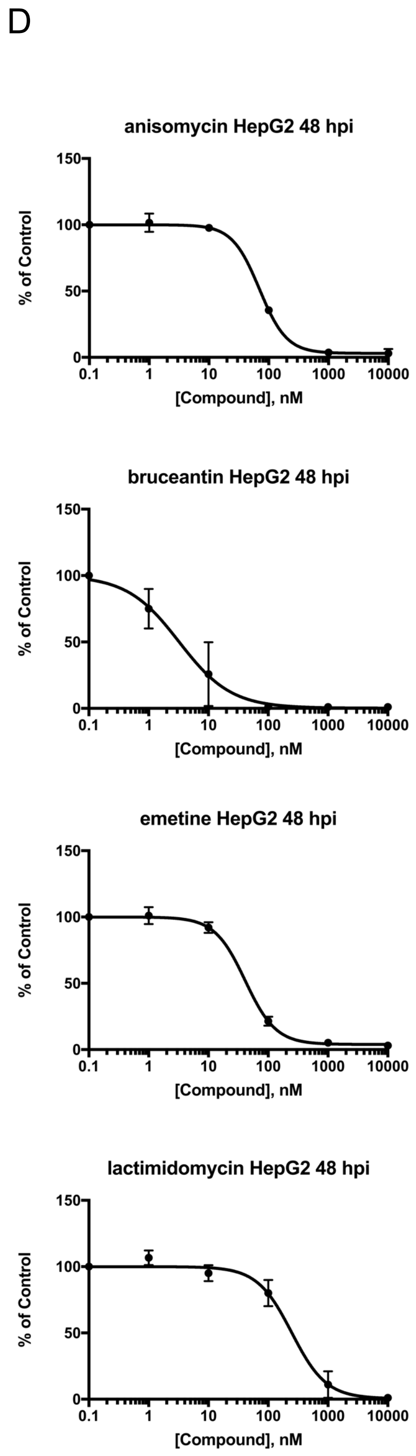


**Figure S10. Estimating potency of translation inhibitors.** Translation was quantified following acute pre-treatment in *P. berghei* liver stages during early schizogony (A) and matching in-image HepG2 cells (B), and in *P. berghei* liver stages during late schizogony (C) and corresponding in-image HepG2 cells (D), using 5-point, 10-fold serial dilution. Plots marked with an asterisk were fit with the bottom of the curve equal to 0, while all others were fit open, as detailed in Methods. Dataset as in Fig. 3 and Fig. S3-2; n ≥ 3 independent experiments.

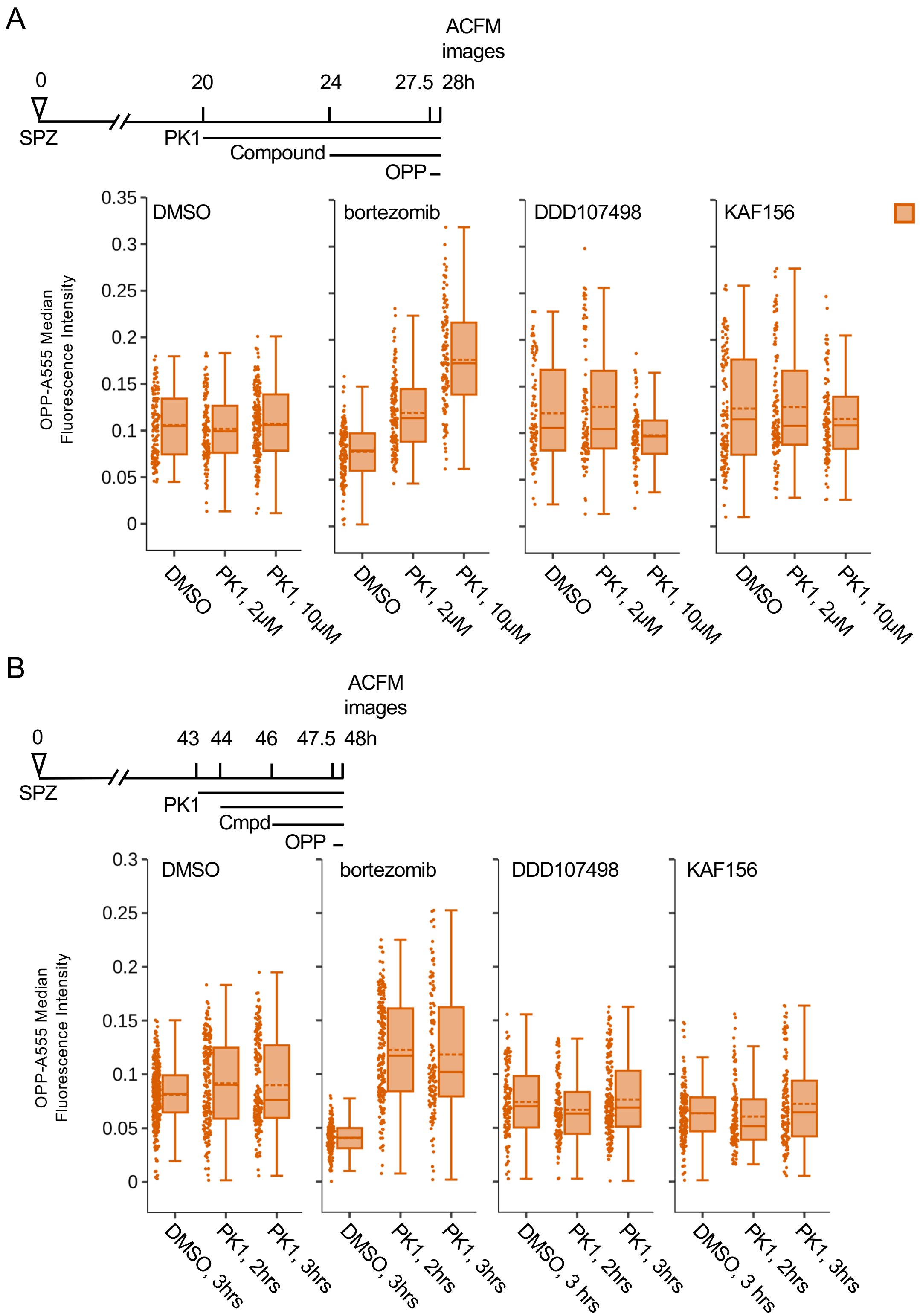


**Figure S11. PK1 pre-treatment modulates bortezomib-induced changes in HepG2 protein synthesis.** Experiment schematics and boxplots quantifying HepG2 translation after PK1 pre-treatment, then addition of DDD107498, KAF156, or bortezomib, as labeled. Each data point represents in-image HepG2 cells corresponding to the single parasite data quantified in Fig. 5A-B. [bortezomib] = 1 μM, [KAF156] = 0.5 μM, and [DDD107498] = 0.1 μM in A-B); [PK1] = labeled in A) and 20 μM in B). Boxplots show cumulative data from n=3 independent experiments, with mean additionally indicated by a dotted line.

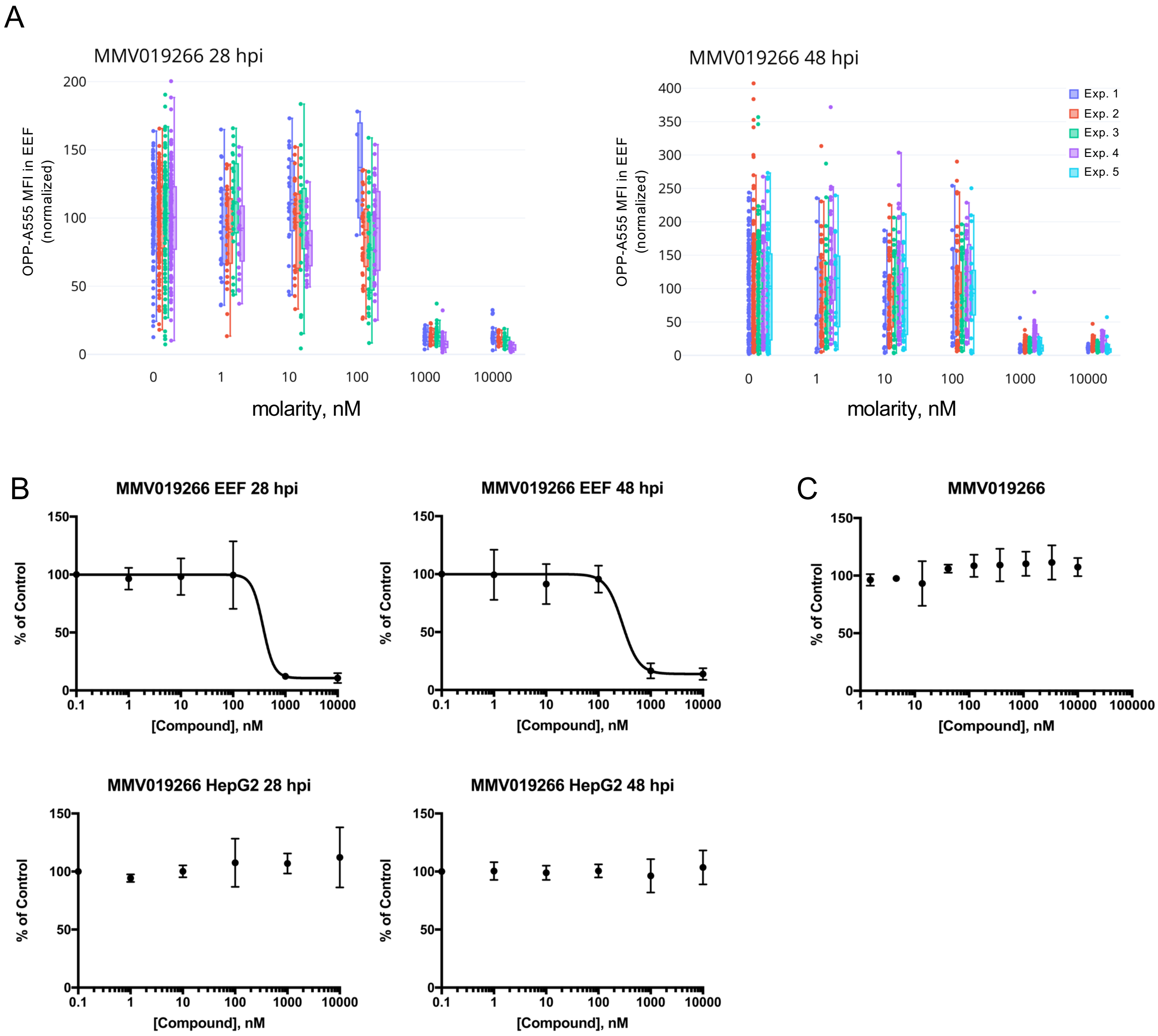


**Figure S12. Concentration-response testing of MMV019266.** Quantification of *P. berghei*-infected HepG2 translation following acute pre-treatment with MMV019266 in concentration response. A) Individual experiments shown for data summarized in Figure 6C, and fitted curves B) for both EEF and HepG2 translation inhibition. C) Fitted HepG2 cytotoxicity concentration response curve following treatment with 10-point, 3-fold serial dilution; maximal concentration 10μM in B-C). All data measured in n≥3 independent experiments.

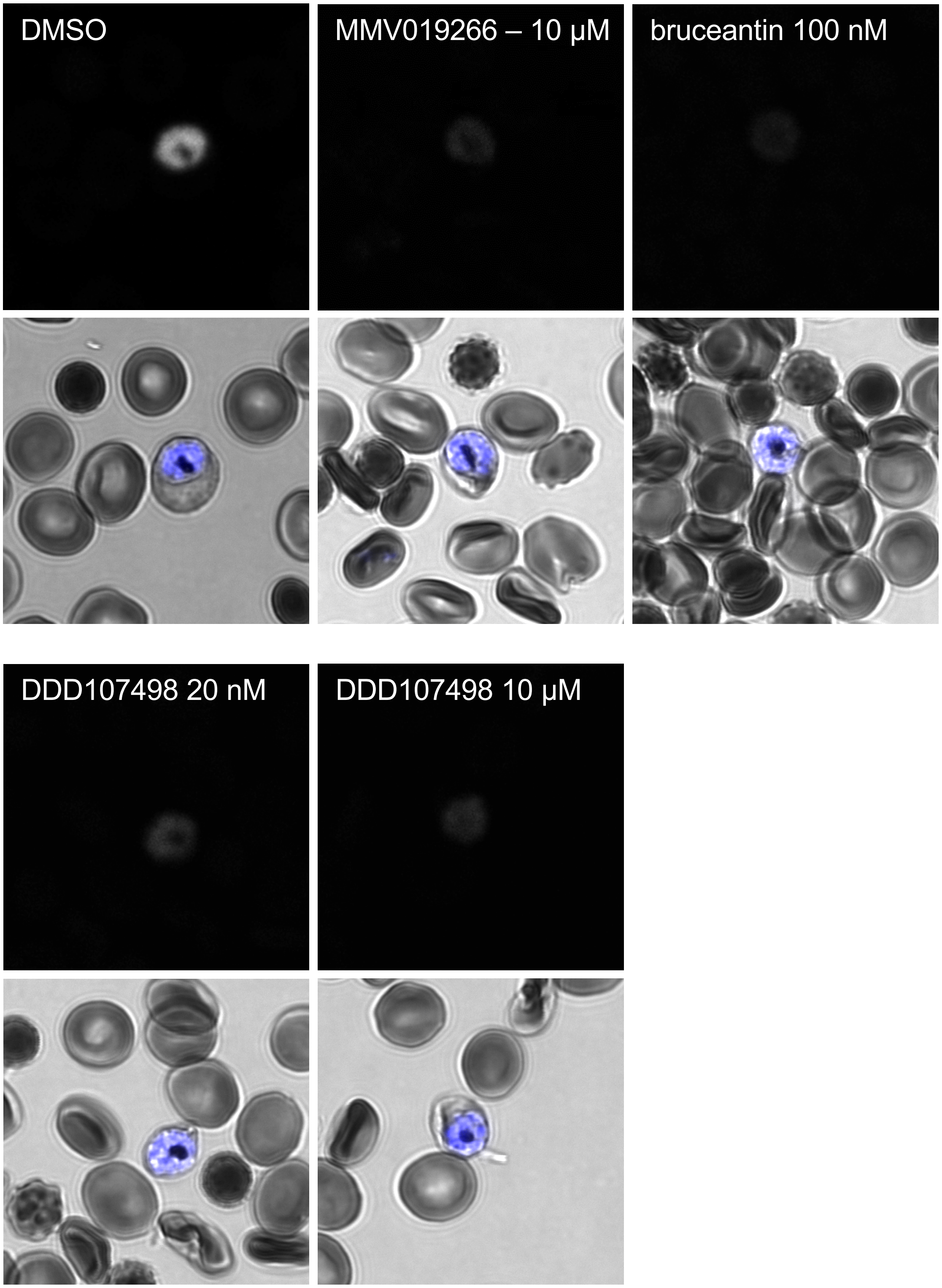


**Figure S13. MMV019266 inhibits *P. falciparum* translation during blood stage schizogony.** Representative single confocal images of *P. falciparum* blood stage schizonts following a 4h acute pre-treatment pulse of compound, as labelled in figure. OPP-A555 labels the nascent proteome (grayscale), and brightfield images are merged with Hoechst-labeled parasite DNA (blue). All images were acquired with identical settings.
